# Spatial microRNA profiling at single-cell resolution by in situ barcoded extension

**DOI:** 10.64898/2026.08.12.744364

**Authors:** Agustín Robles-Remacho, Yimin Zou, Augusta Jensen, Christos Tricopoulos, Marco Grillo, Mats Nilsson

## Abstract

The spatial organization of post-transcriptional regulation is a fundamental yet difficult to access layer of tissue biology. MicroRNAs (miRNAs) are small RNAs with a key role in post-transcriptional regulation, but their short length has excluded them from spatial profiling technologies, leaving them largely unexplored in spatial transcriptomics. Here, we introduce miR-Space, a method that converts individual miRNAs into extended, uniquely barcoded molecules directly in tissue, enabling their spatial detection by in situ sequencing. Across 30 mouse and human brain sections, miR-Space enabled highly multiplexed miRNA profiling at single-molecule and single-cell resolution, joint analysis with mRNA, and implementation on the automated Xenium platform. miR-Space resolved major anatomical regions and cell populations from spatial miRNA expression, identified reproducible cell-associated miRNA signatures, and uncovered previously unknown spatial and cellular distributions of multiple miRNAs. Together, these capabilities establish miR-Space as a framework for integrating miRNAs into spatial transcriptomics, enabling spatial miRNomics at anatomical and single-cell resolution.

## Main

Gene expression is regulated within individual cells and shaped by their spatial position and temporal context in the living organism ^1^. This regulation operates through transcriptional, epigenetic, and post-transcriptional mechanisms that determine when, where, and to what extent genes are expressed^2–4^. These layers are inherently spatial: cells interpret regulatory inputs according to their location, lineage identity, local microenvironment, and surrounding tissue architecture ^5,6^. The integration of intrinsic regulatory programs with local signals and tissue architecture drives cell identity, differentiation, tissue maintenance, and responses to injury^7,8^. Spatial position is therefore a key determinant of how gene expression programs are established and maintained within tissues. MicroRNAs (miRNAs) constitute a major class of post-transcriptional regulators of gene expression^9,10^. These small non-coding RNAs, typically 21-22 nucleotides in length, guide Argonaute-containing complexes to target messenger RNAs through sequence complementarity, leading to transcript degradation or translational repression ^11,12^. By acting across multiple targets, miRNAs form regulatory networks that coordinate diverse cellular programs^13^. Their expression varies across developmental stages, cell states, and cellular identities, linking miRNA activity to distinct biological functions ^14,15^.

Despite their central role in gene regulation, miRNAs remain largely inaccessible to spatial omics^16^. Spatial transcriptomic technologies broadly comprise sequencing-based and imaging-based approaches^17^. Sequencing-based methods capture RNA in tissue and identify transcripts by *ex situ* sequencing, whereas imaging-based methods detect individual targets directly in tissue through *in situ* hybridization and optical decoding^18,19^. Most sequencing-based platforms rely on poly(A) capture and therefore inefficiently recover miRNAs, which naturally lack poly(A) tails. Methods based on *in situ* polyadenylation have expanded spatial profiling to non-polyadenylated RNAs, including miRNAs. Spatial Total RNA (STR) sequencing uses *in situ* polyadenylation to profile coding and noncoding RNA classes^20^, while Patho-DBiT combines *in situ* polyadenylation with microfluidic spatial barcoding in FFPE tissue and has been used to detect region-associated miRNAs^21^. However, neither approach directly measures miRNAs at single-cell resolution: STR profiles spatial capture regions, whereas Patho-DBiT reconstructs single-cell maps through computational integration with histology rather than direct cell assignment. Imaging-based miRNA detection, typically using locked nucleic acid probes^22^ or signal-amplification strategies^23^, has likewise remained limited to individual miRNAs or small target sets^24,25^. Thus, spatial miRNomics still lacks a method that combines highly multiplexed miRNA detection with direct single-cell resolution in intact tissue^26^.

Here, we present miR-Space (spatial microRNA profiling by *in situ* barcoded extension), a strategy for highly multiplexed spatial miRNA detection in intact tissue. miR-Space extends endogenous miRNAs with sequence-encoded barcodes that can be read by in situ sequencing^27,28^, making them compatible with spatial detection workflows designed for longer RNA molecules. We systematically evaluated assay specificity, robustness, and scalability across barcode architectures, experimental conditions, and panel sizes, and further assessed compatibility with joint miRNA–mRNA detection and automated Xenium analysis. In the adult mouse brain, miR-Space resolved regional organization and enabled single-cell analysis of miRNA expression across major cellular populations. In the human cerebral cortex, it identified distinct miRNA distributions across anatomical compartments.

## Results

### miR-Space extends miRNAs in situ with unique barcodes for spatial profiling

miR-Space operates directly in tissue through three main stages. First, following fixation, a long oligonucleotide complementary to the target miRNA, link A, hybridizes to the miRNA and generates an overhang that recruits a second oligonucleotide, link B, carrying a miRNA-specific barcode. Link B is covalently ligated to the miRNA, after which link A is removed, leaving an extended, barcoded miRNA at its original spatial location (**Fig. 1a**). The barcode sequence is designed to be orthogonal to endogenous RNAs, providing a more specific molecular identity than the native miRNA sequence alone. Second, a padlock probe hybridizes to the extended miRNA, with a footprint spanning both the miRNA and its specific barcode. The padlock probe is circularized by ligation and amplified by rolling circle amplification (RCA) to generate a localized amplicon. Third, the resulting amplicons are decoded by *in situ* sequencing-by-hybridization using bridge probes and fluorescent detection oligonucleotides (**Fig. 1b**). Together, miRNA-specific barcodes, padlock-encoded sequence features, and cyclic imaging create a combinatorial coding scheme that enables highly multiplexed identification of miRNA species directly in tissue and spatial and single-cell profiling of miRNA expression (**Fig. 1c-d**).

**Fig. 1.**
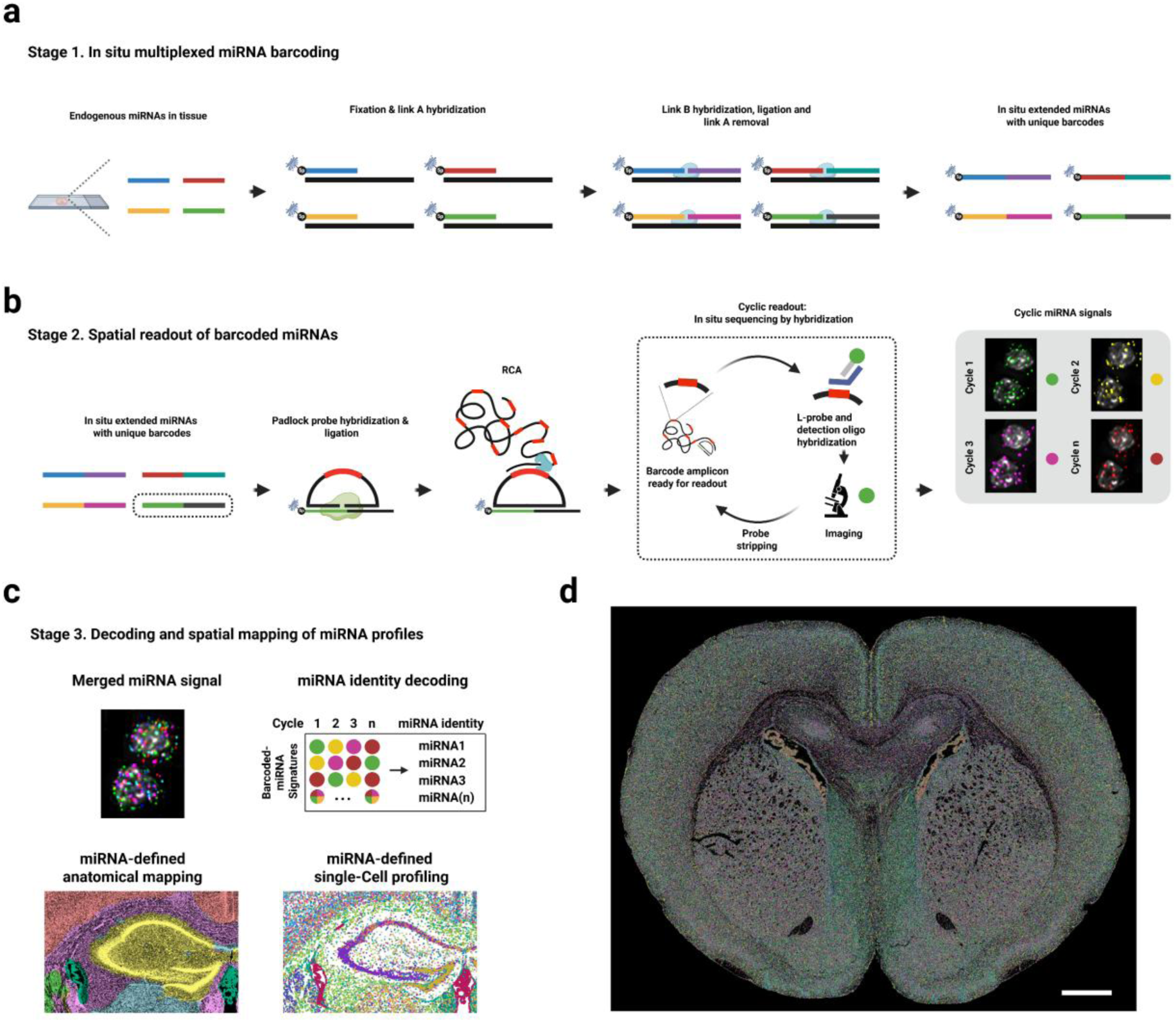
miR-Space extends miRNAs in situ with unique barcodes for spatial profiling. a, Following tissue fixation, endogenous miRNAs are hybridized by link A, which recruits link B carrying a miRNA-specific barcode. Link B is ligated to the target miRNA, and removal of link A leaves an extended, uniquely barcoded miRNA at its original tissue location. b, Barcoded miRNAs are detected by padlock-probe ligation, rolling circle amplification, and cyclic *in situ* sequencing-by-hybridization. c, Combinatorial decoding enables highly multiplexed identification of individual miRNA species for spatial and single-cell profiling. d, Representative spatial map of multiplexed miRNA detection in a mouse brain section. Scale bar, 1 mm.

Target specificity is achieved through two complementary design features. First, each miRNA is extended with an orthogonal barcode designed not to hybridize to endogenous RNAs. Second, the ligation site of the padlock detection probe is positioned within a target-specific sequence. This site can lie within the native miRNA, within the added barcode, or across the miRNA-barcode junction. In the current design, ligation occurs within a miRNA-specific sequence near this junction, while the padlock probe spans both the endogenous miRNA and the added barcode. This architecture enables the use of longer padlock probes with improved GC content and hybridization properties and provides greater flexibility in selecting a unique ligation site. For miRNAs whose full sequence is also present within longer endogenous transcripts (Supporting Fig. 1), barcode extension creates a distinct molecular signature that distinguishes the extended miRNA from these transcripts and reduces cross-reactivity. Multiplexing is achieved through two levels of barcoding: the first barcode is introduced during miRNA extension and defines target identity, whereas the second is encoded in the padlock probe. Together, these design features enable efficient hybridization, specific ligation, and balanced detection across highly multiplexed panels.

### miR-Space enables specific and highly multiplexed spatial miRNA profiling with flexible barcode architectures and no-extension controls

To evaluate the specificity and multiplexing capacity of miR-Space, we designed a nominal 50-plex panel, referred to as version 1, for fresh-frozen adult mouse brain sections. The panel comprised 53 miRNA targets selected based on expression in the mouse brain according to the miRNA Tissue Atlas 2025^29^. miR-Space yielded abundant multiplexed miRNA detections directly in tissue while preserving spatial information (**Fig. 2a**). We next tested whether these signals specifically depended on miRNA extension. No-extension controls were performed on consecutive tissue sections using padlock probes designed to recognize the extended product. Control conditions included omission of both linkers, addition of link B alone, and mismatched linkers, thereby testing whether signal generation required successful extension or could result from nonspecific padlock-probe hybridization to endogenous RNAs or miRNA precursors. Signal levels were markedly reduced under all control conditions, demonstrating that efficient detection required formation of the intended extended miRNA and therefore the presence of a mature miRNA with an accessible 3′-OH end (**Fig. 2b**). A small residual signal remained under no-extension conditions, most prominently for miR-124-3p (**Fig. 2b**). Residual signal levels correlated with those in the complete reaction, consistent with low-level recognition of unextended miRNAs (Supplementary Fig. 2).

**Fig. 2.**
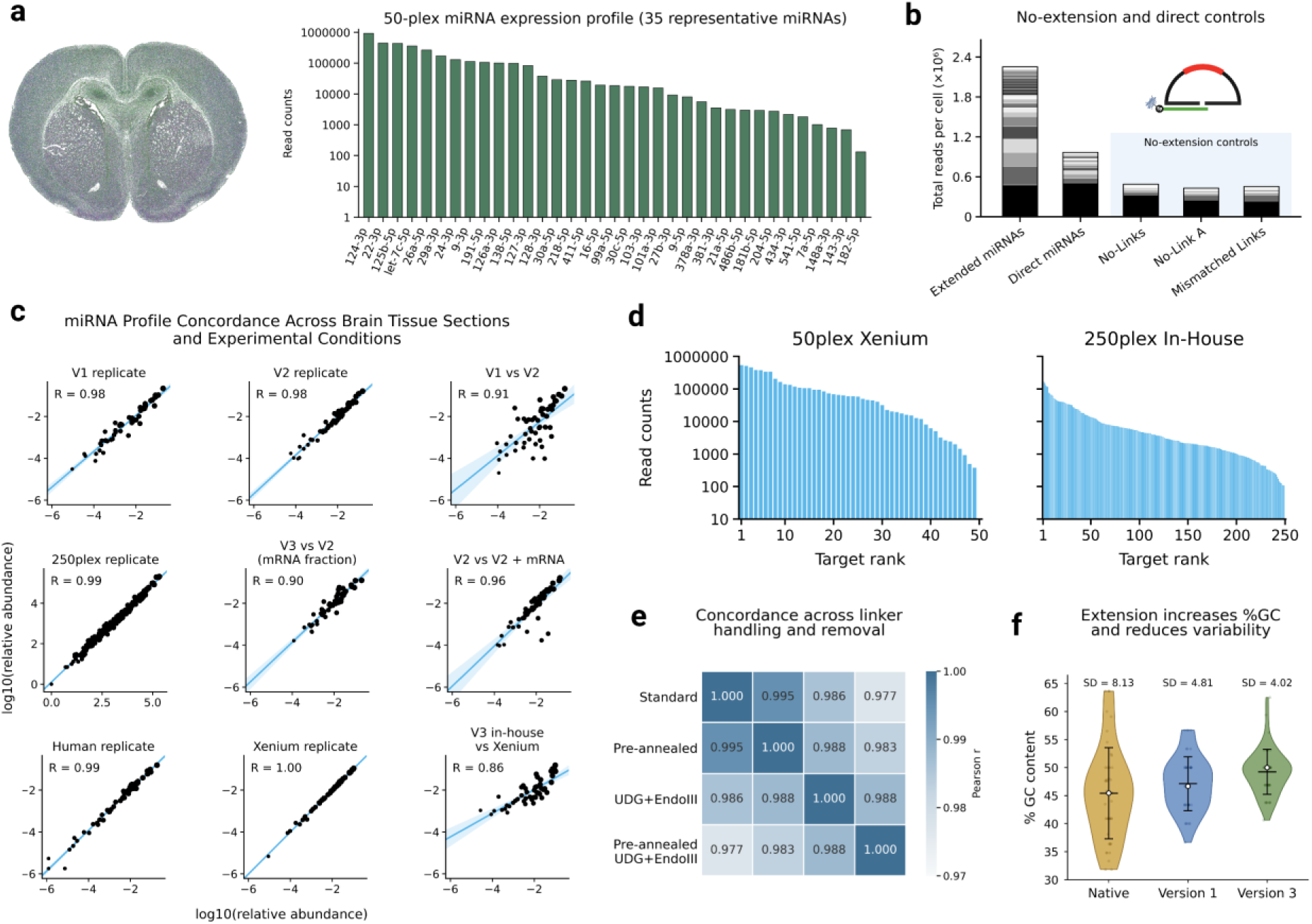
miR-Space enables reproducible, extension-dependent, and scalable spatial miRNA profiling. a, Representative spatial map and expression profile of the version 1 50-plex panel in an adult mouse brain section. Bars show read counts for 35 representative miRNAs. b, Total reads normalized per cell across five consecutive brain sections comparing extended-miRNA detection, direct native-miRNA detection, and extension-specific controls: no linkers, link B only, and mismatched linkers. Grey shades represent individual miRNAs, whereas black denotes miR-124-3p, highlighting its predominance in the control profiles. c, Concordance of miRNA expression profiles across 12 tissue sections from 6 animals spanning panel versions, experimental formats, and platforms. Pearson correlation coefficients are shown. d, Ranked target counts for the version 3 50-plex panel analyzed on Xenium and the 250-plex panel analyzed using the in-house workflow. e, Correlation matrix showing concordance of miRNA profiles across linker-handling and long-linker-removal conditions in four consecutive brain sections. f, GC-content distributions of native miRNAs and extension designs from versions 1 and 3, showing that the longer extension used in version 3 increased GC content and reduced sequence variability.

We next assessed whether miR-Space improved miRNA detection compared with direct targeting of native miRNAs. To this end, we designed a second 50-plex padlock-probe panel containing the same 53 miRNA targets as panel version 1 but targeting only the native miRNA sequences, and compared both approaches in consecutive tissue sections. Direct detection yielded fewer and less balanced signals (Fig. 2b), with most detections concentrated in miR-124-3p and several miRNA spatial patterns remaining undetected. This reduced target coverage and signal diversity limited reliable single-cell analysis. A detailed comparison with direct detection is provided in Supplementary Fig. 2.

We then examined the reproducibility of miR-Space and the influence of barcode identity on the resulting spatial profiles. We designed a second 50-plex panel, referred to as version 2, in which the same miRNAs were assigned a different set of barcodes. We compared replicates within the same panel version and across panel versions, animals, and experimental batches. Spatial miRNA distributions were highly consistent, and the relative abundance of each miRNA within the total miRNA signal was reproducible, with Pearson correlation coefficients above 0.9 (**Fig. 2c**). Additional comparisons between panel versions 1 and 2, including reads detected per cell and ligases tested for compatibility with miR-Space, are provided in Supplementary Fig. 3.

To evaluate scalability and compatibility with mRNA detection, we designed an expanded 250-target panel comprising 259 targets in total, including 208 miRNAs and 51 mRNAs. The assay generated abundant reads across targets (**Fig. 2d**) while preserving the spatial organization of both RNA classes. The corresponding no-extension control is shown in Supplementary Fig. 2. Because Xenium requires longer target sequences for panel design, we developed a third panel, version 3, with an extended target-sequence footprint and miRNAs selected from the expanded in-house panel based on their distinct spatial patterns. These targets were implemented in a nominal 300-plex Xenium panel comprising 50 miRNAs and 247 mRNAs. miR-Space generated robust signals and recovered spatial miRNA distributions on Xenium (**Fig. 2d**). In 12 tissue sections from six animals and one human donor, miRNA expression profiles remained highly concordant between panel versions 1, 2, and 3, replicate 250-target experiments, replicate human sections, and version 3 analyzed using both Xenium and a non-automated in-house workflow in which imaging and decoding were performed independently of the Xenium platform (**Fig. 2c**). Together, these results show that miR-Space generates reproducible spatial miRNA profiles that are robust to barcode reassignment, experimental variation, panel scaling, and platform implementation, while enabling highly multiplexed joint spatial profiling of miRNAs and mRNAs.

miR-Space uses padlock probes that span both the native miRNA and the added barcode sequence, requiring removal of the long miRNA-complementary linkers used during extension. We therefore evaluated a modified workflow incorporating linker pre-annealing and UDG/EndoIII-mediated cleavage, referred to as miR-Space USER, in combination with formamide washing. miRNA profiles remained highly concordant across all conditions (**Fig. 2e**). miR-Space USER produced a modest improvement in signal performance while preserving these profiles (Supplementary Fig. 4); however, given the simplicity of the workflow, formamide treatment alone was used in subsequent experiments. Extension increased and homogenized the GC content of the probe-hybridization regions compared with direct miRNA detection, while the longer target-sequence footprint in version 3 further reduced GC-content variability, enabling more balanced padlock-probe design and detection across the panel (**Fig. 2f**). We then compared miRNA abundance measured by miR-Space Xenium with three independent mouse brain small RNA-seq datasets. All three comparisons showed positive correlations, with only minor biases and no strong abundance-dependent distortion, indicating that miR-Space Xenium preserves relative mature miRNA abundance across the targeted panel (Supplementary Fig. 5). A comparison of miR-Space with current spatial miRNA profiling methods is provided in Supplementary Table 4. Across the linker and padlock-probe libraries, we compiled 1,965 linker and detection-probe sequences targeting 337 miRNAs and 50 mRNAs; the complete sequences and annotations are provided in Supplementary Table 1-3. Code and analysis scripts used in this study are described in the Code Availability section.

### Spatial miRNA profiling resolves anatomical organization in the adult mouse brain

We next assessed whether miR-Space could recover adult mouse brain anatomy from spatial miRNA expression. We analyzed five sections from five mice using the 50-plex v3 panel (three coronal and two sagittal; adult, P28, and P16 samples) and two additional coronal sections using the 250-plex v2 panel. All samples were analyzed using Xenium, except one sagittal section and both 250-plex coronal sections, which were analyzed using the in-house workflow. The resulting multiplexed maps followed recognizable brain architecture and resolved major structures, including cortical, hippocampal, striatal, ventricular, vascular, olfactory, and cerebellar compartments (**Fig. 3a**). Higher-magnification views further revealed distinct local organization within the hippocampus, dentate gyrus, corpus callosum, olfactory bulb, and cerebellar layers. Individual miRNAs showed diverse anatomical distributions, ranging from broad expression to enrichment in specific compartments; representative compartment-enriched examples are shown in **Fig. 3b**. These patterns were reproducible across samples, with no notable differences between developmental stages.

**Fig. 3.**
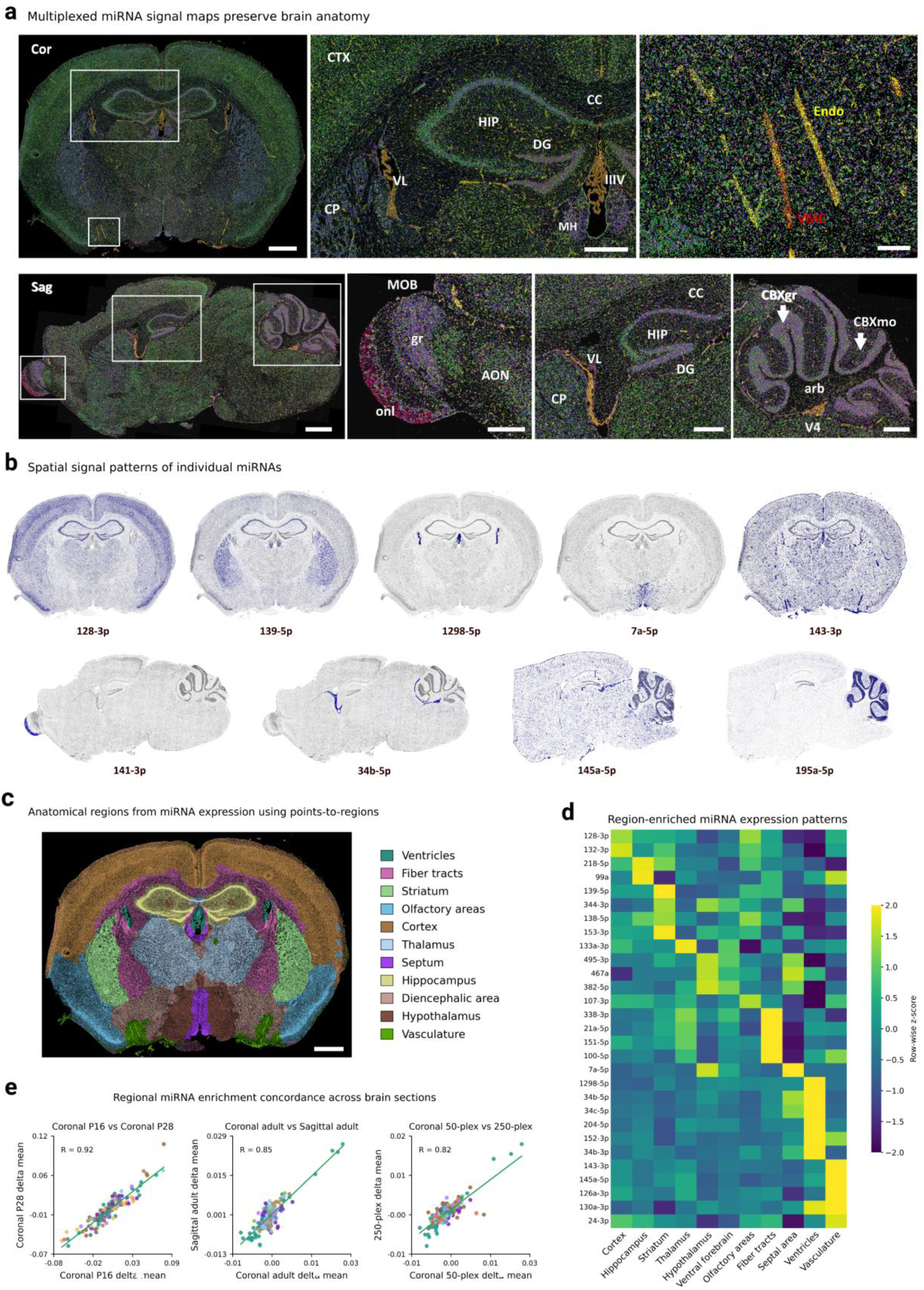
Spatial miRNA expression resolves anatomical organization in the adult mouse brain. a, Multiplexed miRNA maps from coronal (Cor) and sagittal (Sag) mouse brain sections analyzed with the 50-plex version 3 panel, with higher-magnification views of selected anatomical and vascular regions. b, Representative spatial expression patterns of individual miRNAs in coronal and sagittal brain sections, showing enrichment of miR-128-3p in the cortex, miR-139-5p in the striatum, miR-1298-5p and miR-34b-5p in the ventricles, miR-143-3p and miR-145a-5p in the vasculature, miR-7a-5p in the hypothalamus, miR-141-3p in the olfactory bulb, and miR-195a-5p in the cerebellum. c, Anatomical regions identified from miRNA expression using Points2Regions analysis. d, Heatmap of region-enriched miRNA expression derived from the Points2Regions analysis. e, Comparison of regional miRNA enrichment signatures between P16 and P28 coronal sections, adult coronal and sagittal sections, and the Xenium 50-plex and in-house 250-plex panels. Pearson correlation coefficients are shown. Scale bars, 1 mm in full coronal and sagittal views; 500 µm in higher-magnification views, except Endo and VMC, 125 µm. Abbreviations: CTX, cortex; HIP, hippocampus; DG, dentate gyrus; CC, corpus callosum; MH, medial habenula; VL, lateral ventricle; V4, fourth ventricle; CP, choroid plexus; AON, anterior olfactory nucleus; MOB, main olfactory bulb; onl, olfactory nerve layer; arb, arbor vitae; CBXgr, cerebellar granular layer; CBXmo, cerebellar molecular layer; Endo, endothelial cells; VMC, vascular mural cells.

The observed distributions were consistent with established findings from previous studies, including enrichment of miR-128-3p and miR-132-3p in the cortex^30,31^, miR-126a-3p and miR-143-3p in vascular structures^32,33^, miR-204-5p in the choroid plexus^34^, miR-7a-5p in the hypothalamus^35^, miR-141-3p in the olfactory bulb^36^, and miR-195a-5p in the cerebellum^30^. In contrast, several patterns had not been described in the adult mouse brain, including miR-139-5p and miR-153-3p enrichment in the striatum, miR-182-5p in piriform areas, miR-1298-5p and miR-34b/c-5p in ventricular and choroid plexus-associated regions, and miR-495-3p and miR-335-5p in ventral diencephalic regions. These distributions were consistent across barcode versions 1, 2, and 3 and were strongly reduced in no-extension controls, supporting their specificity. Together, these findings show that miR-Space captures established anatomical miRNA organization while revealing previously unknown spatial patterns in the adult mouse brain. Additional spatial patterns of individual miRNAs are shown in Supplementary Fig. 6, with comparison to findings reported in the literature summarized in Supplementary Table 5. Interactive resources for exploring miRNA distributions in coronal and sagittal brain sections are provided in Supplementary Data 1 and 2.

We analyzed anatomical organization using Points2Regions analysis^37,38^, grouping detected molecules into spatially defined territories based on local miRNA enrichment. These territories were anatomically annotated using the Allen Mouse Brain Atlas^39^ as ventricles, fiber tracts, striatum, olfactory areas, cortex, thalamus, septum, hippocampus, diencephalic areas, hypothalamus, and vasculature, with spatial distributions consistent with established brain anatomy (**Fig. 3c**). We then generated a row-wise normalized heatmap showing the predominant miRNA enrichment within each territory, linking individual miRNAs to specific anatomical regions (**Fig. 3d**). Regional miRNA signatures, quantified as delta mean values, were subsequently compared across independent animals, developmental stages, sectioning planes, and panel configurations. Coronal sections from P16 and P28 mice showed highly correlated regional profiles, with a Pearson correlation coefficient of 0.92 (**Fig. 3e**). Comparisons of adult sections with P16 and P28 showed similarly strong concordance (Supplementary Fig. 7). Regional signatures were also preserved between coronal and sagittal sections, with a Pearson correlation coefficient of 0.85 (**Fig. 3e**). Scaling miR-Space from the Xenium 50-plex panel to the in-house 250-plex panel preserved regional miRNA organization, with a Pearson correlation of 0.82 across the two experimental platforms (**Fig. 3e**). Together, these results demonstrate robust preservation of regional miRNA signatures across animals, developmental stages, sectioning planes, platforms, and highly multiplexed panels.

### Spatial single-cell miRNA profiling resolves cell-type identity in intact tissue

Next, we evaluated miR-Space at single-cell resolution across six coronal brain sections from five mice. Four sections were analyzed on Xenium using a 300-plex panel comprising 50 miRNAs and the predefined 247-mRNA Mouse Brain Gene Expression Panel, including two consecutive adult sections from the same animal and one section each from P28 and P16 mice. Two additional sections from independent mice were analyzed using the in-house workflow with a 250-target panel. Following cell segmentation, miRNA molecules were decoded and assigned to individual cells to generate single-cell miRNA expression matrices. Cell populations were defined exclusively from their miRNA profiles using Scanpy, and differential expression was assessed using the Wilcoxon rank-sum test. Clusters were initially interpreted from their spatial distribution, with annotations subsequently supported by established mRNA markers co-expressed within the same cells (**Fig. 4a**). Across the six sections, the analysis encompassed 676,863 segmented cells.

**Fig. 4.**
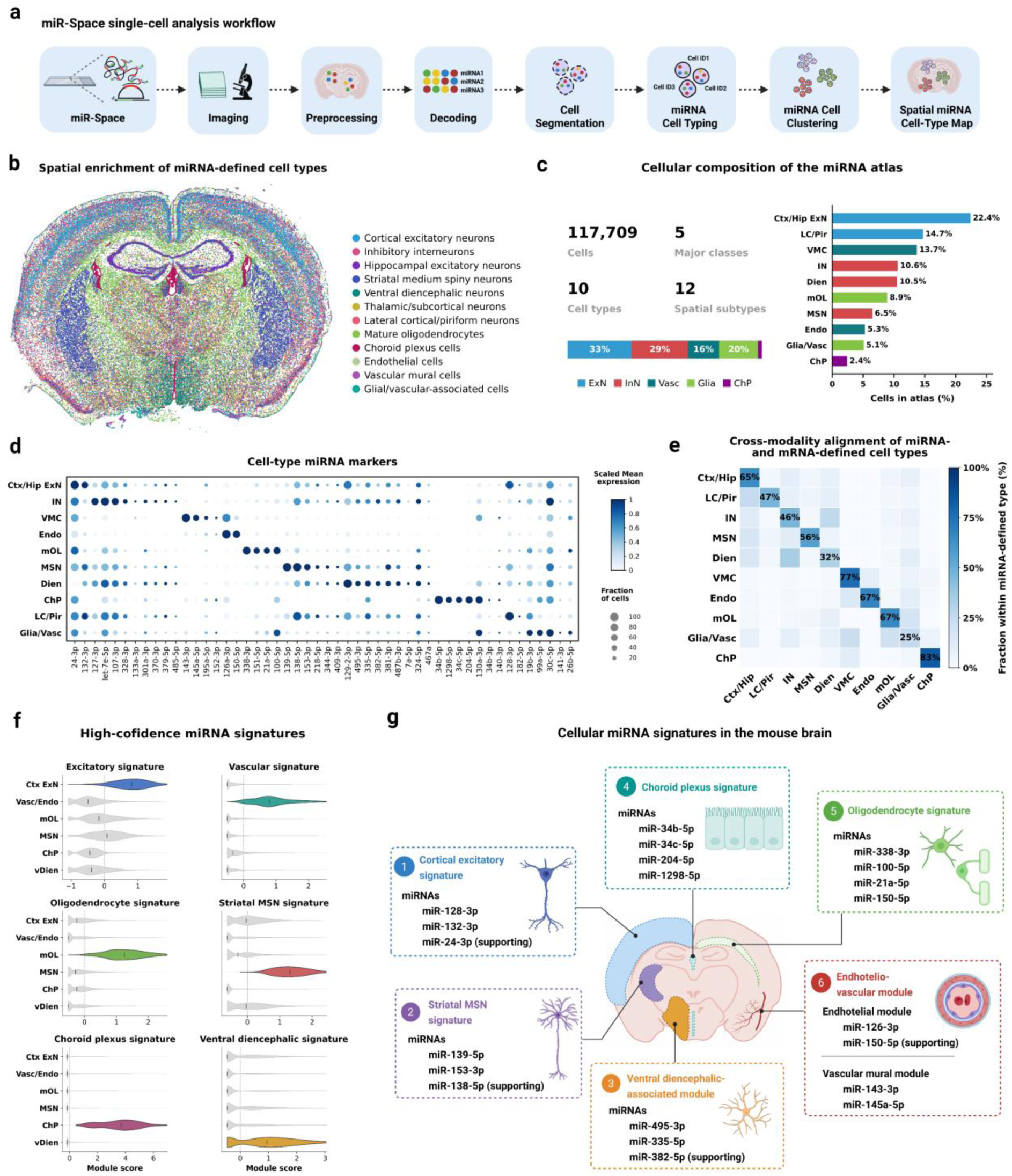
Single-cell spatial miRNA profiling recovers major cell classes in the adult mouse brain. a, Overview of the miR-Space single-cell analysis workflow, including imaging, preprocessing, miRNA decoding, cell segmentation, miRNA-based cell typing, clustering, and spatial mapping. b, Spatial distribution of miRNA-defined cell types across a representative coronal mouse brain section. c, Cellular composition of the miRNA atlas, including total cell number, major classes, cell types, spatial subtypes, and relative population abundance. d, Dot plot of miRNA markers across identified cell types; dot size indicates the fraction of expressing cells and color indicates scaled mean expression. e, Cross-modality comparison between miRNA-defined and mRNA-defined cell types, shown as the fraction of cells within each miRNA-defined population assigned to each mRNA-defined cell type. f, Module scores for six high-confidence cell-type-associated miRNA signatures across major cell populations. g, Spatial summary of the six miRNA signatures and their representative miRNAs across the mouse brain. Abbreviations: ChP, choroid plexus; Ctx/Hip ExN, cortical and hippocampal excitatory neurons; Dien, diencephalic neurons; Endo, endothelial cells; Glia/Vasc, glial and vascular-associated cells; IN, inhibitory neurons; LC/Pir, lateral cortical and piriform populations; mOL, mature oligodendrocytes; MSN, medium spiny neurons; VMC, vascular mural cells.

To characterize the cellular organization in greater detail, we analysed a representative adult coronal section. miR-Space identified 117,709 cells encompassing ten cell types, which could be grouped into five major classes and further resolved into twelve spatial subtypes (**Fig. 4b,c**). Neuronal populations included cortical and cortico-hippocampal excitatory neurons, lateral cortical/piriform neurons, striatal medium spiny neurons (MSNs), inhibitory neurons, ventral diencephalic neurons, and thalamic/hypothalamic neurons. Non-neuronal populations comprised mature oligodendrocytes, glial/vascular-associated cells, endothelial cells, vascular mural cells, and choroid plexus cells. Several populations were further resolved into spatial subtypes occupying distinct anatomical territories. Each cell type exhibited a distinct miRNA expression profile, defined by scaled mean expression and the fraction of miRNA-positive cells (**Fig. 4d**). Additional characterization, including UMAPs, reads and features per cell, and spatial distributions of individual cell types is provided in Supplementary Fig. 7.

Using the same adult coronal section, we independently classified the 117,709 cells using the 259-plex mRNA expression matrix. We then compared the identity assigned to each cell based on mRNA with that obtained from the 50-plex miRNA panel. The two classifications showed the strongest agreement for well-defined populations, including excitatory neurons, medium spiny neurons, endothelial cells, mature oligodendrocytes, and choroid plexus cells, whereas agreement was lower for more heterogeneous or regionally defined populations, such as lateral cortical/piriform and glial/vascular-associated groups (**Fig. 4d**). The ventral diencephalic territory also showed differences, with the miRNA-based analysis identifying a ventral diencephalic population distinct from that defined by the mRNA-based classification, as discussed below. Differences likely reflect the greater number of markers and optimization of the mRNA panel for cell-type classification, while miRNA profiles provide additional biological information not fully captured by the measured mRNA fraction alone. We then systematically evaluated all annotated populations in the adult coronal section by integrating statistical enrichment, marker specificity within the corresponding cellular or anatomical compartment, spatial localization, and concordance with mRNA-defined cell identities. The single-cell analyses supporting population assignments and associated miRNA markers are provided in Supplementary Tables 5-7. This analysis identified six high-confidence cell-associated miRNA modules, defined as miRNA signatures associated with a specific cell population. These signatures corresponded to cortical excitatory neurons, vascular and endothelial cells, mature oligodendrocytes, striatal MSNs, choroid plexus/ependymal cells, and ventral diencephalic neurons. For each module, miRNA expression was z-score standardized across all selected cells, and a per-cell module score was calculated as the mean z-score of its constituent miRNAs. Comparison of module scores across annotated populations showed preferential enrichment of each module in its corresponding target population, with substantially lower scores in non-target populations (**Fig. 4f**).

We next assessed the robustness and scalability of these signatures in the adjacent adult section, the P16 and P28 samples, and independent in-house 250-plex replicates. The same six miRNA modules consistently distinguished their corresponding cell populations across all samples, demonstrating reproducibility across biological specimens and developmental stages, while remaining preserved in the high-plex configuration encompassing 209 miRNAs and thereby supporting the scalability of the approach (Supplementary Figs. 9 and 10). Together, these six high-confidence modules defined distinct and reproducible cell-associated miRNA signatures across neuronal, glial, and vascular populations, revealing a structured layer of cell-type-associated miRNA expression in the mouse brain (**Fig. 4g**). Several of the cell-type signatures were consistent with previously established associations, including miR-128-3p and miR-132-3p in cortical excitatory neurons^40^, miR-338-3p and miR-100-5p in mature oligodendrocytes^41^, miR-143-3p and miR-145a-5p in vascular mural cells^33^, and miR-126a-3p in endothelial cells^32^. The analysis also identified previously uncharacterized miRNA signatures, including miR-139-5p, miR-138-5p, and miR-153-3p to striatal medium spiny neurons, miR-335-5p and miR-495-3p to ventral diencephalic neurons, and the miR-34b-5p/miR-34c-5p/miR-204-5p/miR-1298-5p signature to choroid plexus/ependymal cells.

Examination of the mRNA content of cells comprising the six high-confidence miRNA modules showed canonical markers for their corresponding cell populations, supporting concordance between miRNA- and mRNA-based cell identities (Supplementary Table 6). In the ventral diencephalon, the miRNA-defined population was characterized by the inhibitory neuronal markers Gad2, Cdh13, Rasgrf2, and Rab3b, together with enrichment of miR-335-5p and miR-495-3p. By contrast, when cells were classified using the mRNA-only expression matrix, the ventral diencephalic population showed Slc17a6 expression, lacked the Gad2-associated profile, and did not show enrichment of miR-335-5p or miR-495-3p. Both miRNAs have been linked to neuronal excitability, synaptic plasticity, activity-dependent regulation, and neuronal differentiation^42,43^, supporting a potential role in region-specific neuronal programs. Together, these results show that the miRNA-defined cellular signature in the ventral diencephalon differs from that obtained by mRNA-based classification, resolving two distinct cellular populations within the same anatomical region and highlighting molecular information captured by miRNA profiling that is not reflected by mRNA expression alone.

### miR-Space enables spatial miRNA profiling in human tissue

To evaluate the applicability of miR-Space to human specimens, we applied a 50-plex miRNA panel to sections of fresh-frozen human cerebral cortex. The panel was designed using miRNAs reported in the miRNATissueAtlas 2025^29^. miR-Space generated abundant spatially resolved signals and revealed distinct expression patterns across the tissue, with clear differences between cortical gray matter, underlying white matter, and pial-vascular compartments (**Fig. 5a,b**). A points-to-region analysis was then used to define these three anatomical regions and compare their miRNA profiles. The most enriched miRNAs in each compartment, as well as the overall regional enrichment profiles, were highly correlated between replicate sections (**Fig. 5c**). Individual miRNAs showed preferential localization to distinct anatomical compartments, including miR-127-3p, miR-128-3p, and miR-22-3p in cortical gray matter, miR-181a-5p in white matter, and miR-143-3p and miR-451a in the pial-vascular compartment (**Fig. 5d,e**), consistent with previous studies reporting region- and cell-type-associated miRNA expression in cortical, vascular smooth muscle, and erythroid populations ^33,44,45^. A row-wise normalized heatmap identified the miRNAs most strongly enriched in each anatomical compartment, while regional dominance analysis highlighted those contributing most prominently to the molecular identity of each region (**Fig. 5e,f**). Together, these results demonstrate that miR-Space can resolve reproducible and anatomically organized miRNA expression patterns in human tissue, recovering regional signatures consistent with known brain and vascular miRNA biology. A summary of all 30 tissue sections analyzed in this study is provided in Supplementary Table 8.

**Fig. 5.**
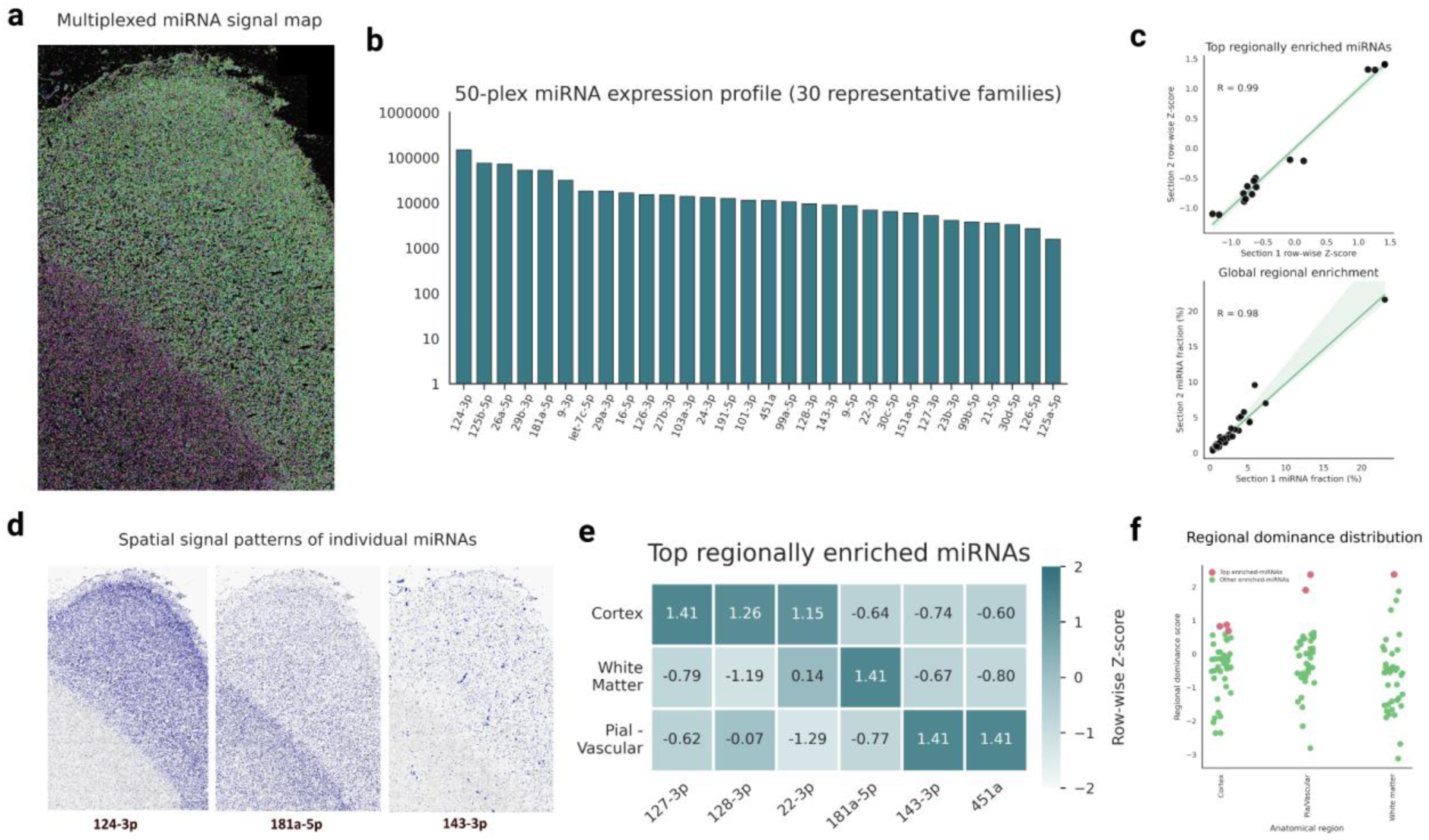
Spatial miRNA profiling in the human cerebral cortex with miR-Space. a, Multiplexed spatial map of miRNA signals detected with a 50-plex panel in a human cerebral cortex section. b, Total expression profile of 30 representative miRNA families, shown on a logarithmic scale. c, Reproducibility between replicate sections based on points-to-region analysis, comparing the enrichment scores of the top regionally enriched miRNAs (top) and the complete regional enrichment profiles (bottom). Pearson correlation coefficients are indicated. d, Representative spatial distributions of individual miRNAs preferentially localized to cortical gray matter, white matter, or the pial-vascular compartment. e, Row-wise normalized heatmap derived from the points-to-region analysis, showing the miRNAs most strongly enriched across the three anatomical compartments. f, Regional dominance analysis showing the contribution of individual miRNAs to the molecular profile of each anatomical region; the most regionally enriched miRNAs are highlighted.

## Discussion

miR-Space addresses a persistent technological gap in spatial transcriptomics by enabling highly multiplexed miRNA detection at anatomical and single-cell resolution in intact tissue. miRNAs are difficult targets for spatial profiling because of their short length, limited sequence space, and variable GC content. The main advantage of miR-Space is that it does not rely on direct detection of the native miRNA sequence alone. Instead, endogenous miRNAs are extended in situ with target-specific barcodes, generating longer molecules that are more compatible with spatial transcriptomics platforms. This reduces constraints imposed by the native miRNA sequence, increases flexibility in probe design, and provides an additional sequence feature for target identification. The strong reduction in signal observed in no-extension controls, together with the agreement obtained across independent barcode configurations, supports that the spatial profiles recovered by miR-Space depend on formation of the intended extended product rather than on nonspecific recognition of endogenous RNA.

An important feature of the approach is its scalability and compatibility with existing spatial workflows. The same biological profiles were preserved across different barcode designs, experimental batches, panel configurations, and both in-house and Xenium implementations. In addition, scaling from the initial 50-plex configuration to panels containing more than 200 miRNAs did not disrupt the regional or cell-associated signatures identified with the smaller panel. This suggests that the extension strategy is not restricted to a particular probe architecture or decoding system and can be adapted as spatial platforms evolve. The implementation on Xenium is particularly relevant because it shows that short RNA species can be incorporated into automated imaging-based spatial workflows that were originally designed for longer RNA targets. More generally, in situ barcode extension may provide a useful strategy for adapting other short or sequence-constrained RNA species to spatial transcriptomic technologies.

Beyond the technical development, the data support the existence of a structured spatial layer of miRNA expression in the brain. Regional miRNA profiles followed anatomical organization and were preserved across independent animals, developmental stages, sectioning planes, panel configurations, and across mouse and human tissue. This reproducibility indicates that miRNA distributions are not simply driven by local differences in total RNA abundance or assay performance, but reflect stable features of tissue organization. Several of the recovered patterns were consistent with previous studies, whereas others had not been spatially characterized in the adult mouse brain. Importantly, miR-Space also revealed previously uncharacterized cell-associated miRNA signatures, including signatures linked to striatal medium spiny neurons, ventral diencephalic neurons, and choroid plexus/ependymal populations. These observations suggest that spatial miRNA profiling can identify regulatory signatures that are not apparent from existing anatomical or transcriptomic descriptions and provide candidates for further functional investigation.

The single-cell analysis further indicates that miRNA expression contains information related to cellular identity. Major neuronal, glial, and vascular populations could be distinguished using miRNA expression alone, and several high-confidence miRNA modules remained associated with the same populations across biological replicates and expanded high-plex experiments. Importantly, however, miRNA- and mRNA-based classifications were not fully equivalent. For well-defined populations, the two modalities showed strong agreement, whereas lower correspondence was observed for more heterogeneous or regionally defined groups. The ventral diencephalic region was particularly informative, as the miRNA-based analysis identified a population with an inhibitory-associated mRNA profile and enrichment of miR-335-5p and miR-495-3p that differed from the population obtained using the mRNA-only classification. This difference support that miRNA expression can capture molecular information that is not fully represented by the measured mRNA fraction.

This complementary information may be one of the main biological advantages of spatial miRNA profiling. miRNAs act at the post-transcriptional level and therefore represent a regulatory layer distinct from transcript abundance itself. Cells with broadly similar transcriptional identities may differ in miRNA programs associated with activity, differentiation, stress responses, or local physiological context^46,47^. In this setting, miRNAs are unlikely to replace mRNA markers for broad cell classification, but they may refine the molecular characterization of populations that are transcriptionally similar or anatomically heterogeneous. Integrating spatial miRNA measurements with mRNAs, proteins, and other molecular layers could identify cell populations with similar transcriptional profiles but distinct regulatory states, including programs associated with activation, stress, differentiation, or altered physiology. Mapping miRNAs in intact tissue may also clarify their cellular origin and reveal how post-transcriptional programs are organized across anatomical boundaries and cellular niches^48^. This information could help determine how regulatory states are coordinated between neighboring populations and how they contribute to intercellular communication, development, and tissue maturation^49–51^. Such analyses may be particularly informative in heterogeneous tissues, including tumors, where spatially restricted miRNA programs could support biological interpretation and biomarker discovery^52,53^. Spatial miRNomics, integrated with current transcriptomic and proteomic measurements, may therefore provide an additional layer of regulatory information.

Overall, miR-Space provides a framework for integrating miRNAs into spatial transcriptomic analysis. By converting short endogenous miRNAs into extended, barcoded molecules while preserving their original tissue location, the method enables spatial miRNomics to be analyzed alongside established anatomical and single-cell transcriptomic approaches. Joint profiling of miRNA and mRNA expression enables direct investigation of how post-transcriptional regulatory programs are organized across cell types and tissue architecture. The recovery of both established and previously uncharacterized spatial and cell-associated miRNA signatures further indicates that this additional molecular layer captures biologically meaningful organization of post-transcriptional regulation. Together, these findings establish miR-Space as a general strategy for bringing miRNA biology into spatially resolved molecular analysis and for expanding the regulatory information accessible in intact tissues.

## Methods and Materials

### Sample collection

A total of 30 fresh-frozen tissue samples were processed, including 26 mouse brain samples and 4 human cerebral cortex samples. For the in-house experiments, 20 commercially obtained mouse brain sections derived from four different mice and four commercially obtained human cerebral cortex frozen sections derived from the same donor were used. All samples were purchased from Zyagen (San Diego, CA, USA). The mouse samples were derived from 10-week-old adult CD1 (ICR) mice, whereas sex was not recorded. Animals were sourced from Charles River (Wilmington, MA, USA), and tissue collection was performed on the day of arrival or after a short acclimation period. Brains were snap-frozen in OCT without fixative. The mouse sections included both coronal and sagittal orientations. The human cerebral cortex frozen sections were derived from a 56-year-old Caucasian male donor and were prepared from freshly harvested tissue snap-frozen in OCT. According to the supplier, all mouse and human sections were cut at a thickness of 7-10 µm. In addition, 6 mouse brain sections used for Xenium analyses were prepared from tissue blocks obtained from the laboratory of Michael Ratz (Karolinska Institutet, Stockholm, Sweden). Three sections were obtained from wild-type C57BL/6 brains from 2-month-old adult mice; these comprised two coronal sections and one sagittal section. The remaining three sections were obtained from wild-type CD-1 brains corresponding to developmental stages of 1 week, 2 weeks, and 28 days. All tissue blocks were prepared by perfusion and freezing prior to transfer and were sectioned for analysis. Tissue blocks were sectioned at a thickness of 10 µm and placed onto Xenium slides. All slides were stored at -80 °C in a dry environment with silica beads until further processing.

### Reagents, materials, and oligonucleotides

The following reagents and materials were used throughout the MiR-Space workflow. SplintR Ligase (25,000 U mL^-^¹) and recombinant albumin, molecular biology grade, were obtained from New England Biolabs/NEB. T4 DNA ligase (5 U µL^-^¹), formamide, RNase-free 20× SSC, HCl (approximately 37% in water), ATP solution (100 mM), and SlowFade Gold Antifade Mountant were obtained from Thermo Fisher Scientific. Formaldehyde solution (37%), EDC-HCl powder, and glycerol (>99.5%) were obtained from Merck/Sigma-Aldrich. Molecular biology-grade water, 1-methylimidazole (99%), DAPI, and 24 × 24 mm Menzel coverslips were obtained from VWR. Silica gel beads with indicator were obtained from Merck. PBS, PBS containing 0.05% Tween-20, NaCl (5 M), Tris buffer (1 M), and EDTA (0.5 M) were provided by Karolinska Institutet. Phi29 polymerase was produced by the Karolinska Institutet Protein Science Core Facility (PSF). All on-tissue reactions were carried out using SecureSeal™ hybridization chambers (Grace Bio-Labs, GBL621502; Merck/Sigma-Aldrich). All version 1, 2, and 3 oligonucleotides were obtained from Integrated DNA Technologies (IDT). The linker and padlock-probe libraries comprised 1,965 sequences targeting 337 miRNAs and 50 mRNAs are provided in Extended Data 1 and 2.

### Sample Fixation and Permeabilization

Slides were retrieved from the -80 °C freezer and allowed to air-dry for 15 min in a dry environment with silica beads to prevent water condensation during warming to room temperature. Tissue sections were then fixed with 3.7% formaldehyde for 15 min at room temperature and washed three times with PBS. For EDC-mediated crosslinking, a working solution containing 0.16 M EDC in 0.13 M imidazole buffer (pH 8.0, 0.3 M NaCl) was prepared immediately before use. The 0.13 M imidazole buffer was prepared in H_2_O, adjusted to pH 8.0 with HCl, and NaCl was added to a final concentration of 0.3 M. Freshly prepared 1.6 M EDC stock in RNase-free H_2_O was then added to the buffer to reach a final EDC concentration of 0.16 M. Sections were covered with EDC/imidazole solution and incubated for 2 h, followed by three washes in PBS. Sections were then permeabilized with 0.1 M HCl for 5 min at room temperature, washed three times with PBS, and dehydrated through a graded ethanol series consisting of 70%, 85%, and 100% ethanol for 1 min each. Following fixation, permeabilization, and dehydration, tissue sections were ready for downstream processing with MiR-Space protocol.

### miR-Space protocol

The MiR-Space protocol was carried out over four consecutive days, ending either with global imaging of RCA products or with preparation of the sections for sequencing cycles. Typically, up to two sequencing cycles can be performed per day using a microscope; therefore, additional days may be required depending on the total number of sequencing cycles performed. All on-tissue reactions were carried out using SecureSeal™ hybridization chambers in a total volume of 100 µL. On day 1, sections were washed for 1 min in 0.01% PBS-Tween at room temperature, followed by two washes in PBS, and were kept in PBS until addition of the miRLink probe hybridization mix. To prepare this mixture, miRLink probes were heat-denatured immediately before use and added to a buffer containing 2× SSC and 20% formamide to a final concentration of 20 nM each. The miRLink probe hybridization mix was added to the hybridization chambers, the chambers were sealed, and the slides were incubated overnight at 37 °C.

On day 2, sections were washed in SSC/formamide wash buffer (2× SSC, 10% formamide), washed twice in PBS, and incubated for 4 h at 22 °C in miRLink ligation mix containing 1× T4 DNA ligase buffer, 5% PEG4000, 1 mM ATP, 0.2 mg mL^-^¹ BSA, and T4 DNA ligase at a final concentration of 0.15 Weiss U µL^-^¹. Sections were then washed twice in PBS, and the hybridization chambers were replaced with new ones. The long miRLink was then stripped by washing the sections three times in 2× SSC, followed by three washes in 100% formamide for 1 min each, and a further three washes in 2× SSC. The sections were kept in 2× SSC until the padlock probe hybridization mix was added. miRNA and/or mRNA padlock probes were heat-denatured immediately before use and added to a buffer containing 2× SSC and 20% formamide to a final concentration of 20 nM each. The resulting padlock probe hybridization mix was applied to the sections, the chambers were sealed, and the slides were incubated overnight at 37 °C.

On day 3, sections were washed twice for 1 min in SSC/formamide wash buffer (2× SSC, 10% formamide), washed twice in PBS, and incubated for 4 h at 37 °C in padlock probe ligation mix containing 1× SplintR ligase reaction buffer and SplintR ligase at a final concentration of 0.5 U µL^-^¹. Following two PBS washes, RCA was carried out overnight at 30 °C in a mixture containing 1× Phi29 polymerase buffer, 5% glycerol, 0.25 mM dNTPs, 0.2 mg mL^-^¹ BSA, an initiation primer at a final concentration of 200 nM, and Phi29 polymerase at a final concentration of 1 U µL^-^¹. On day 4, sections were processed either for global visualization of RCA products or for initiation of sequencing cycle 1. For global visualization, sections were incubated with an AF488-labelled anchor detection oligonucleotide to label all RCA products and were then subjected to imaging. Before sequencing cycle 1, anchor detection oligonucleotides were stripped by washing the sections three times in 2× SSC, followed by three washes in 100% formamide for 1 min each and a further three washes in 2× SSC. Sections not used for anchor-based visualization were processed directly after RCA.

Sequencing cycling was implemented through iterative rounds of hybridization of barcode-specific L-probes, followed by hybridization of fluorophore-labelled detection oligonucleotides, imaging, signal stripping and re-hybridization. For sequencing cycle 1, sections were washed once in PBS-Tween and once in PBS, followed by hybridization of L-probes (200 nM) for 1 h at room temperature in hybridization buffer containing 1× SSC and 20% formamide. After a further wash in PBS-T and PBS, detection oligonucleotides labelled with distinct fluorophores (AF750, Cy5, Cy3, AF488 and ATTO 425; 200 nM each) were hybridized together with DAPI for 1 h at room temperature in the same buffer. Sections were then washed three times in PBS-T, mounted and imaged. Sequencing was performed through iterative cycles of L-probe and fluorophore-labelled detection oligonucleotide hybridization, imaging, signal removal and re-hybridization. For subsequent sequencing cycles, coverslips were removed by gentle incubation in PBS until detachment. Slides were then washed twice to remove residual mounting medium and air-dried. L-probes and fluorophore-labelled detection oligonucleotides were subsequently stripped by three washes in 2× SSC, followed by three washes in 100% formamide 1 min each, and a further three washes in 2× SSC, before proceeding to the next round of L-probe hybridization.

### miR-Space adaptation to the Xenium workflow and oligonucleotide panels

The MiR-Space protocol was adapted by incorporating the pre-hybridization steps into the Xenium *In Situ* Gene Expression v1 workflow. Tissue sections were placed onto Xenium Slides (10x Genomics Xenium Slides & Sample Prep Reagents, PN-1000460) and stored at -80 °C. As described in Section 2.3, slides were removed from -80 °C, air-dried under dry conditions at room temperature, and subsequently subjected to fixation and permeabilization as previously described. Cassette assembly was then performed according to the manufacturer’s instructions, and all on-slide reactions were carried out in a total volume of 500 µL. Slides were then washed for 1 min in 0.01% PBS-Tween at room temperature, followed by two washes in PBS, and kept in PBS until the miRLink probe hybridization mix was added. The miRLink probe hybridization mix was prepared as described in Section 2.4, with the final volume adjusted to 500 µL for use in the Xenium slide. Next, the lyophilized pre-designed panel probes and custom panel probes were resuspended in TE buffer, pH 8.0, vortexed twice for 15 s, incubated at room temperature for 5 min, and briefly centrifuged before being added to the probe hybridization mix. Probe hybridization and all subsequent steps were then carried out according to the instructions provided in the Xenium In Situ Gene Expression v1 workflow. Incubations were performed using a Xenium Thermocycler Adaptor (PN-3000954) in a Biometra PCR Thermal Cycler (Analytik Jena, Germany). After completion of the wet-lab workflow, samples were processed on a Xenium Analyzer (PN-1000529). For Xenium analyses, the Xenium Mouse Brain Gene Expression Panel (247 genes; 1000462) and the Add-on Custom 50 Gene Panel (1000652) were used.

### Imaging

Image acquisition was performed on a Leica DMi8 inverted epifluorescence microscope equipped with a Lumencor SPECTRA X external LED illumination source, an LMT200-HS automated multi-slide stage, and a Leica DFC9000 GTC sCMOS camera. All images were acquired using an HC PL APO 20×/0.80 objective with a numerical aperture of 0.80. Multispectral acquisition was performed using the DFT51011 filter set, consisting of a filter cube and an external emission filter wheel, for spectral separation. Images were acquired in six fluorescence channels corresponding to Alexa 750 (810 nm), Cy5 (683 nm), Cy3 (595 nm), Alexa 488 (515 nm), DAPI (432 nm), and AT425 (480 nm). Regions of interest were defined using LAS X software (v3.7.5.24914). Z-stacks spanning 10 µm were acquired at 0.5 µm intervals. Raw and processed image data were stored in SciLifeLab FAIR Storage (Storage3), which provides backup and controlled access for authorized project users.

### Image preprocessing, decoding and cell segmentation

Raw images were preprocessed using an in-house pipeline available in Moldia/ISS2025 GitHub repository. Images were organized by imaging cycle, region, tile and channel, deconvolved using RedLionFish with a synthetic point-spread function and 50 iterations, and converted into maximum-intensity projections. These were then wrapped into OME-TIFF files, aligned across cycles and stitched using Ashlar across fields of view with 10% overlap, with the nuclear channel used as the registration reference. Stitched mosaics were finally re-tiled into smaller images for downstream analysis. Decoding was performed using an in-house pipeline available in the Moldia/ISS2025 GitHub repository. Preprocessed re-tiled images were converted to SpaceTx format together with the corresponding codebook, registered across cycles using DAPI as the reference channel, and decoded with a Starfish-based workflow using PRMC mode after MH normalization. Spots were detected with an intensity threshold of 0.002 and sigma values ranging from 1 to 10 over 30 steps. Decoded reads were then quality-filtered by retaining only reads with a minimum per-cycle quality score above 0.4. The final output was a decoded CSV file containing the spatial coordinates and gene identity of each decoded signal. Cell segmentation was performed with Cellpose using the stitched DAPI image from the first imaging cycle (channel 4). Diameter was estimated automatically, and segmented nuclei were expanded by 20 pixels to approximate cell boundaries. The resulting segmentation masks were used for downstream spatial assignment of decoded signals. Tissue domains were manually annotated using the corresponding mouse coronal or sagittal images from the Allen Brain Atlas as anatomical reference.

### Point-to-region analysis and anatomical annotation

Point-to-region analysis was performed using Points2Regions on the spatial coordinates and target identities of quality-filtered miRNA molecules. For the coronal section, spatial binning was performed using a pixel width of 28, a minimum of eight molecules per pixel, and a smoothing parameter of 9, followed by clustering into 25 spatial regions. For the sagittal section, miRNAs represented by fewer than 350 decoded molecules were excluded, and the analysis was performed using a pixel width of 75, a minimum of five molecules per pixel, a smoothing parameter of 8, and 20 spatial clusters. The resulting clusters were manually assigned to anatomical compartments according to their spatial distribution and the corresponding reference sections from the Allen Brain Atlas. Region-associated miRNAs were identified using group-versus-rest comparisons and differences in mean expression between each region and the remaining tissue. Regional miRNA profiles were summarized using mean expression and the fraction of spatial units containing each miRNA. Enrichment was evaluated by comparing each annotated anatomical region against all remaining regions. Regional patterns were compared between coronal and sagittal sections using shared miRNAs and harmonized anatomical compartments. Heat maps were generated from standardized regional expression values, and selected region-associated miRNAs were ordered according to the region in which they showed the highest relative enrichment.

### Xenium data import and AnnData generation

Xenium output files were imported and processed to generate cell-by-feature matrices containing expression counts, cell metadata, and spatial coordinates. Feature identifiers corresponding to the custom miRNA probes were replaced with their associated miRNA names. Features were then classified as miRNAs or mRNAs, and separate AnnData objects were generated for the miRNA-only, mRNA-only, and combined miRNA-mRNA datasets. Cell boundaries and transcript-to-cell assignments were obtained from the standard Xenium output. In contrast, for datasets generated using the custom imaging workflow, decoded molecules were assigned to Cellpose-derived cell masks according to their spatial coordinates.

### Single-cell quality control, normalization and dimensionality reduction

Quality-control metrics were calculated separately for miRNA and mRNA features. For both the mRNA- and miRNA-based analyses, cells containing fewer than five total detected transcripts were excluded. Features detected in fewer than 30 cells were removed before downstream analysis. Raw counts were retained in a separate AnnData layer. Expression matrices were normalized by total counts per cell using the median library size as the normalization target and were subsequently log-transformed using log1p. No feature scaling or highly variable feature selection was applied, and all retained targets from each panel were used for dimensionality reduction. Dimensionality reduction and clustering were performed independently for the miRNA and mRNA expression matrices using Scanpy. Principal-component analysis was calculated using the ARPACK solver, with 11 components for the miRNA analysis and 10 components for the mRNA analysis. Neighborhood graphs were constructed using 30 neighbors, the first 10 principal components, and correlation distance. UMAP embeddings were generated using a minimum distance of 0.15, a spread of 1.0, and a random seed of 0. Leiden clustering was performed using the igraph implementation with two iterations and a random seed of 0, using resolutions of 1.5 for the miRNA analysis and 0.6 for the mRNA analysis.

### Cell clustering, annotation and differential expression

miRNA-based clusters were defined exclusively from the miRNA expression matrix. Cluster identities were initially interpreted from their spatial distribution and miRNA marker profiles and were subsequently supported by established mRNA markers detected in the same cells. Independent mRNA-based cell identities were assigned using the mRNA expression matrix, established marker genes, spatial distribution, and manual curation. Anatomical localization was assessed using the corresponding coronal or sagittal sections from the Allen Brain Atlas as reference. Cluster-associated miRNAs and mRNAs were identified by group-versus-rest differential expression analysis using the Wilcoxon rank-sum test. Groups containing fewer than ten cells were excluded from differential ranking. Reproducibility was evaluated across the four mouse brain sections analyzed at single-cell resolution. Cell-type composition was calculated as the proportion of retained cells assigned to each harmonized cell class. For each sample and cell type, miRNA profiles were summarized using mean log-transformed expression and, where indicated, the fraction of cells expressing each miRNA. Pairwise concordance between sections was assessed using Pearson correlation coefficients for both cell-type composition and cell-type-specific miRNA expression profiles. Only cell types and miRNAs shared between the compared datasets were included. miRNA- and mRNA-based cell classifications were generated independently and compared at the level of individual cells using their shared Xenium cell identifiers. Closely related clusters were harmonized into broader cell classes before comparison. Agreement between classifications was summarized using cell-by-cell contingency tables and the proportion of cells assigned to corresponding miRNA- and mRNA-defined populations. Concordance was evaluated separately for well-defined cell types and for heterogeneous or regionally defined populations.

### Statistics

Expression profiles were compared across datasets after normalization to the same number of cells. Normalized miRNA counts were converted to relative frequencies within each dataset. For comparisons between panel or barcode versions, replicate-specific frequencies were averaged to generate mean version-level profiles. Pearson correlation coefficients were calculated from paired measurements of shared miRNAs. Negative-control comparisons were expressed as log2 fold changes calculated from normalized miRNA abundance in the complete reaction relative to the corresponding control condition. Total abundance was reported both as normalized reads per dataset and as reads per cell. Unless otherwise stated, statistical analyses and graphical representations were generated using Python and GraphPad Prism.

## Supplementary Information

- Supplementary Figures 1–10 are provided in the Supplementary Information file.
- Supplementary Tables 1–3 are provided as separate files.
- Supplementary Tables 4–8 are provided in the Supplementary Information file.
- Supplementary Data 1 and 2 comprise interactive tissue maps of representative miR-Space coronal and sagittal mouse brain sections and are available at the following links:

https://serve.scilifelab.se/projects/mir-space-xenium-wrd/

https://serve.scilifelab.se/projects/mir-space-xenium-sagital-itq/

## Code availability

The custom code used to generate and analyze the results presented in this study is available to editors and reviewers during peer review via the GitHub repository https://github.com/Moldia/miR-Space. Upon publication, the code will be made publicly available on GitHub and archived in Zenodo with a permanent DOI.

## Acknowledgements

A.R.R. thanks the Marie Skłodowska-Curie Actions for support from a Postdoctoral Fellowship (Project 101209017, Spatial miRNomics) and the Fundación Alfonso Martín Escudero for support from a postdoctoral fellowship. This work was supported by project grants from Cancerfonden (reference number 24-3457), the Swedish Research Council (reference number 2024-02533), and the the Science for Life Laboratory (SciLifeLab) Technology Development Project (TDP) “Spatial miRNomics” (2026–2027). We thank the In Situ Sequencing Facility at Stockholm University/SciLifeLab for their advice and technical recommendations. We thank Michael Ratz and Jasmin Leibrock (Karolinska Institutet) for providing the mouse brain samples used in Xenium.

## References

1. Liu, L. et al. Spatiotemporal omics for biology and medicine. Cell vol. 187 4488–4519 Preprint at 10.1016/j.cell.2024.07.040 (2024).

2. Klemm, S. L., Shipony, Z. & Greenleaf, W. J. Chromatin accessibility and the regulatory epigenome. Nature Reviews Genetics vol. 20 207–220 Preprint at 10.1038/s41576-018-0089-8 (2019).

3. Brito Querido, J., Díaz-López, I. & Ramakrishnan, V. The molecular basis of translation initiation and its regulation in eukaryotes. Nature Reviews Molecular Cell Biology vol. 25 168–186 Preprint at 10.1038/s41580-023-00624-9 (2024).

4. Cramer, P. Organization and regulation of gene transcription. Nature vol. 573 45–54 Preprint at 10.1038/s41586-019-1517-4 (2019).

5. Collinet, C. & Lecuit, T. Programmed and self-organized flow of information during morphogenesis. Nature Reviews Molecular Cell Biology vol. 22 245–265 Preprint at 10.1038/s41580-020-00318-6 (2021).

6. Armingol, E., Officer, A., Harismendy, O. & Lewis, N. E. Deciphering cell–cell interactions and communication from gene expression. Nature Reviews Genetics vol. 22 71–88 Preprint at 10.1038/s41576-020-00292-x (2021).

7. Hannezo, E. & Heisenberg, C. P. Mechanochemical Feedback Loops in Development and Disease. Cell vol. 178 12–25 Preprint at 10.1016/j.cell.2019.05.052 (2019).

8. Clevers, H., Loh, K. M. & Nusse, R. An Integral Program for Tissue Renewal and Regeneration: Wnt Signaling and Stem Cell Control. http://science.sciencemag.org/.

9. Ambros, V. Minireview MicroRNAs: Tiny Regulators with Great Potential. Cell vol. 107 (2001).

10. Bartel, D. P. Metazoan MicroRNAs. Cell vol. 173 20–51 Preprint at 10.1016/j.cell.2018.03.006 (2018).

11. Shang, R., Lee, S., Senavirathne, G. & Lai, E. C. microRNAs in action: biogenesis, function and regulation. Nature Reviews Genetics vol. 24 816–833 Preprint at 10.1038/s41576-023-00611-y (2023).

12. Bartel, D. P. MicroRNAs: genomics, biogenesis, mechanism, and function. Cell 116(**2**), 281–97 (2004).

13. Bracken, C. P., Scott, H. S. & Goodall, G. J. A network-biology perspective of microRNA function and dysfunction in cancer. Nature Reviews Genetics vol. 17 719–732 Preprint at 10.1038/nrg.2016.134 (2016).

14. Ludwig, N. et al. Distribution of miRNA expression across human tissues. Nucleic Acids Res. 44, 3865–3877 (2016).

15. De Rie, D. et al. An integrated expression atlas of miRNAs and their promoters in human and mouse. Nat. Biotechnol. 35, 872–878 (2017).

16. Robles-Remacho, A., Zou, Y., Grillo, M. & Nilsson, M. Spatially resolved microRNA expression in tissues: technologies, challenges, and opportunities. Trends in Genetics vol. 41 1131–1143 Preprint at 10.1016/j.tig.2025.06.005 (2025).

17. Moses, L. & Pachter, L. Museum of spatial transcriptomics. Nat. Methods 19, 534–546 (2022).

18. Rao, A., Barkley, D., França, G. S. & Yanai, I. Exploring tissue architecture using spatial transcriptomics. Nature 596, 211–220 (2021).

19. Vickovic, S. et al. Visualization and analysis of gene expression in tissue sections by spatial transcriptomics. Science (1979). 353, 78–82 (2016).

20. McKellar, D. W. et al. Spatial mapping of the total transcriptome by in situ polyadenylation. Nat. Biotechnol. (2022).

21. Bai, Z. et al. Spatially exploring RNA biology in archival formalin-fixed paraffin-embedded tissues. Cell 187, 6760–6779.e24 (2024).

22. Pena, J. T. G. et al. miRNA in situ hybridization in formaldehyde and EDC - Fixed tissues. Nat. Methods 6, 139–141 (2009).

23. Gan, J. et al. MicroRNA-375 restrains the progression of lung squamous cell carcinoma by modulating the ERK pathway via UBE3A-mediated DUSP1 degradation. Cell Death Discov. 9, (2023).

24. Nagarajan, M. B., Tentori, A. M., Zhang, W. C., Slack, F. J. & Doyle, P. S. Spatially resolved and multiplexed MicroRNA quantification from tissue using nanoliter well arrays. Microsyst. Nanoeng. 6, (2020).

25. Ji, X. et al. Sequencing-free spatial profiling of post-transcriptional regulation in fresh tissues using nanoneedle arrays. *Nat*. Biomed. Eng. 10.1038/s41551-026-01745-0 (2026) doi:10.1038/s41551-026-01745-0.

26. Robles-Remacho, A. & Nilsson, M. Spatial miRNomics: towards the integration of microRNAs in spatial biology. Nat. Rev. Genet. 10.1038/s41576-025-00819-0 (2025) doi:10.1038/s41576-025-00819-0.

27. Ke, R. et al. In situ sequencing for RNA analysis in preserved tissue and cells. Nat. Methods 10, 857–860 (2013).

28. Gyllborg, D. et al. Hybridization-based in situ sequencing (HybISS) for spatially resolved transcriptomics in human and mouse brain tissue. Nucleic Acids Res. 48, E112 (2020).

29. Keller, A. et al. MiRNATissueAtlas2: An update to the human miRNA tissue atlas. Nucleic Acids Res. 50, D211–D221 (2022).

30. Bak, M. et al. MicroRNA expression in the adult mouse central nervous system. RNA 14, 432–444 (2008).

31. Mellios, N. et al. MiR-132, an experience-dependent microRNA, is essential for visual cortex plasticity. Nat. Neurosci. 14, 1240–1242 (2011).

32. Wang, S. et al. The Endothelial-Specific MicroRNA miR-126 Governs Vascular Integrity and Angiogenesis. Dev. Cell 15, 261–271 (2008).

33. Xin, M. et al. MicroRNAs miR-143 and miR-145 modulate cytoskeletal dynamics and responsiveness of smooth muscle cells to injury. Genes Dev. 23, 2166–2178 (2009).

34. Lepko, T. et al. Choroid plexus-derived miR-204 regulates the number of quiescent neural stem cells in the adult brain. EMBO J. 38, (2019).

35. Lee, H.-J., Palkovits, M. & Scott Young III, W. MiR-7b, a MicroRNA up-Regulated in the Hypothalamus after Chronic Hyperosmolar Stimulation, Inhibits Fos Translation. www.ebi. (2006).

36. Choi, P. S. et al. Members of the miRNA-200 Family Regulate Olfactory Neurogenesis. Neuron 57, 41–55 (2008).

37. Andersson, A. et al. Points2Regions: Fast, interactive clustering of imaging-based spatial transcriptomics data. Cytometry Part A 105, 677–687 (2024).

38. Marco Salas, S., et al. Optimizing Xenium In Situ data utility by quality assessment and best-practice analysis workflows. Nat. Methods 22, 813–823 (2025).

39. Wang, Q. et al. The Allen Mouse Brain Common Coordinate Framework: A 3D Reference Atlas. Cell 181, 936–953.e20 (2020).

40. He, M. et al. Cell-Type-Based Analysis of MicroRNA Profiles in the Mouse Brain. Neuron 73, 35–48 (2012).

41. Zhao, X. et al. MicroRNA-Mediated Control of Oligodendrocyte Differentiation. Neuron 65, 612–626 (2010).

42. Heiland, M. et al. MicroRNA-335-5p suppresses voltage-gated sodium channel expression and may be a target for seizure control. Proc. Natl. Acad. Sci. U. S. A. 120, (2023).

43. Inouye, M. O., Colameo, D., Ammann, I., Winterer, J. & Schratt, G. miR-329– and miR-495–mediated Prr7 down-regulation is required for homeostatic synaptic depression in rat hippocampal neurons. Life Sci. Alliance 5, (2022).

44. Wang, W. X., Huang, Q., Hu, Y., Stromberg, A. J. & Nelson, P. T. Patterns of microRNA expression in normal and early Alzheimer’s disease human temporal cortex: White matter versus gray matter. Acta Neuropathol. 121, 193–205 (2011).

45. Yu, D. et al. miR-451 protects against erythroid oxidant stress by repressing 14-3-3ζ. Genes Dev. 24, 1620–1633 (2010).

46. Bahl, E. et al. Using deep learning to quantify neuronal activation from single-cell and spatial transcriptomic data. Nature Communications 15, (2024).

47. Scott, E. Y. et al. Integrating single-cell and spatially resolved transcriptomic strategies to survey the astrocyte response to stroke in male mice. Nature Communications 15, (2024).

48. Ding, D. Y. et al. Quantitative characterization of tissue states using multiomics and ecological spatial analysis. Nat. Genet. 57, 910–921 (2025).

49. Pham, D. et al. Robust mapping of spatiotemporal trajectories and cell–cell interactions in healthy and diseased tissues. Nature Communications 14, (2023).

50. Yao, Z. et al. Spatial transcriptomics reveals coordinated ventricular patterning and maturation in the developing human heart. Nat. Commun. 10.1038/s41467-026-74476-0 (2026) doi:10.1038/s41467-026-74476-0.

51. Lohoff, T. et al. Integration of spatial and single-cell transcriptomic data elucidates mouse organogenesis. Nat. Biotechnol. 40, 74–85 (2022).

52. Barkley, D. et al. Cancer cell states recur across tumor types and form specific interactions with the tumor microenvironment. Nat. Genet. 54, 1192–1201 (2022).

53. Sorin, M. et al. Single-cell spatial landscapes of the lung tumour immune microenvironment. Nature 614, 548–554 (2023).

