## Supplementary Information for "Spatial microRNA profiling at single-cell resolution by in situ barcoded extension"

### Index


Supplementary Figure 1. miRNA extension reduces off-target detection ... 3

Supplementary Figure 2. No-extension and direct controls confirm specificity ... 5

Supplementary Figure 3. miR-Space reproducibility and ligase compatibility ... 7

Supplementary Figure 4. miR-Space USER version for enzymatic linker removal ... 9

Supplementary Figure 5. Crossplatform comparison of miR-Space Xenium with small RNA-seq ... 11

Supplementary Figure 6. Spatial atlas of individual miRNA expression across the mouse brain ... 13

Supplementary Figure 7. Single-cell analysis of mRNA and miRNA expression and candidate miRNA-mRNA relationships ... 15

Supplementary Figure 8. Spatial distribution of miRNA-defined cell types across the mouse brain ... 17

Supplementary Figure 9. Reproducibility and cell-type specificity of high-confidence miRNA signatures across independent datasets ... 19

Supplementary Figure 10. ell-type specificity of high-confidence miRNA modules across annotated adult brain cell types ... 21

Supplementary Table 4. Comparison of miR-Space with spatial transcriptomics technologies ... 23

Supplementary Table 5. Literature-based validation and novelty of miR-Space spatial miRNA patterns ... 25

Supplementary Table 6. Integrated interpretation of miRNA-defined cell populations and associated miRNA signatures ... 29

Supplementary Table 7. Single-cell assessment of miRNA markers ... 32

Supplementary Table 8. Best-supported miRNA markers of high-confidence cell populations ... 37

Supplementary Table 6. Tissue sections used in the development of miR-Space ... 39


Supplementary Table 1-3 are provided in a separate document.

Supplementary Data 1 and 2 are provided as interactive online resources.

#### Supplementary Figure 1. miRNA extension reduces off-target detection

Supplementary Fig. 1 illustrates how miRNA barcode extension reduces potential off-target detection from longer transcripts containing overlapping miRNA sequences. One example is mmu-miR-24-3p, whose full sequence is present within the Aopep mRNA. In Supplementary Fig. 1a, the padlock-probe ligation site lies entirely within the sequence shared with the Aopep transcript, preventing selective discrimination of the mature miRNA from the longer mRNA. Barcode extension shifts the ligation site toward the added barcode sequence (Supplementary Fig. 1b). In this design, the padlock probe spans the miRNA-barcode junction, with the ligation site positioned at the miRNA terminus. For the Aopep transcript, only one arm of the padlock probe can hybridize, preventing productive ligation. A no-extension control further supports the specificity of this strategy, with detectable signal generated only after miRNA extension. This indicates that the probe targeting the extended miRNA does not amplify signal derived from the overlapping Aopep transcript (Supplementary Fig. 1e).

A second example is miR-204-5p, whose sequence partially overlaps with the Fam120b mRNA. In Supplementary Fig. 1c, the original ligation site lies within the region shared with the Fam120b transcript, whereas barcode extension relocates the ligation site outside the overlapping region (Supplementary Fig. 1d). Across probe versions, strong signals were detected following miRNA extension, while no-extension controls showed markedly reduced signal. Together, these results indicate that productive padlock-probe ligation depends on formation of the extended miRNA product.

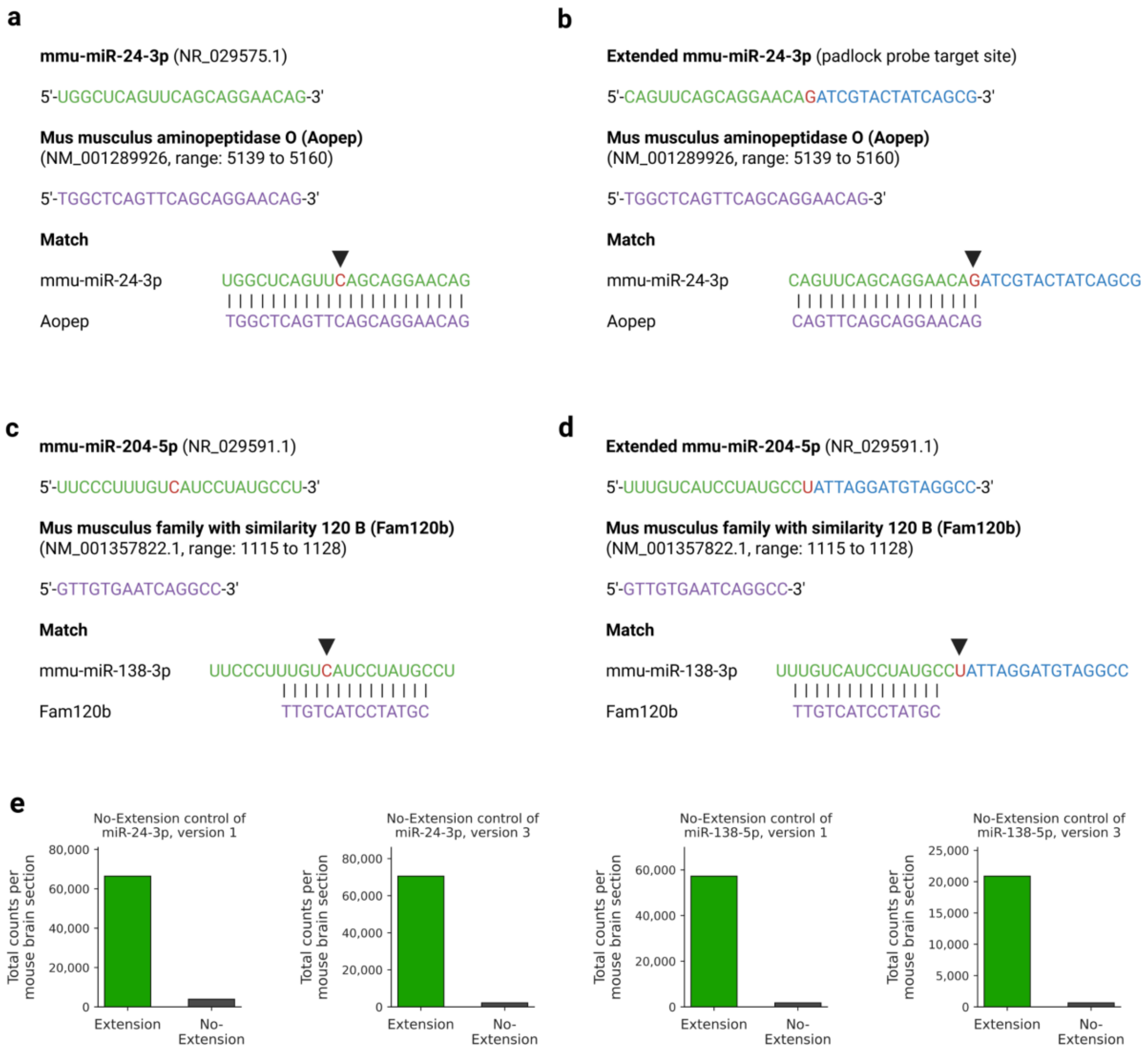


**Supplementary Fig. 1 | miRNA extension reduces off-target detection.**

a, Sequence overlap between miR-24-3p and Aopep mRNA before extension. The ligation site is necessarily contained within the overlapping Aopep mRNA sequence and is indicated by an arrow. b, Barcode extension shifts the padlock-probe ligation site, indicated by an arrow, toward the added barcode. As a result, part of the padlock probe cannot hybridize to Aopep, preventing productive ligation on the mRNA. c, Partial sequence overlap between miR-204-5p and Fam120b mRNA before extension. d, Extension relocates the ligation site outside the shared sequence, thereby preventing productive ligation on Fam120b. e, No-extension controls targeting miR-24-3p and miR-138-5p. Extension produced substantially higher counts than the corresponding no-extension conditions, confirming that the detected signal originates from the extended product and that the padlock probe alone does not generate nonspecific signal.

#### Supplementary Figure 2. No-extension and direct controls confirm specificity

Supplementary Fig. 2 provides a detailed analysis of the no-extension controls introduced in Fig. 2 and evaluates whether miR-Space signal depends on successful formation of the extended miRNA target. In Supplementary Fig. 2a-c, the complete extension reaction was compared with conditions lacking both linkers, containing linker B only, or using target-mismatched linker pairs. In all cases, disruption of linker-mediated extension substantially reduced miRNA detection relative to the complete, correctly matched reaction. This reduction was evident both from the positive fold changes observed for individual miRNAs and from the approximately fivefold decrease in total reads per cell. Despite the lower signal, residual expression profiles remained positively correlated with those obtained under complete-extension conditions, suggesting that the remaining reads most likely arise from limited residual hybridization of padlock probes to their intended native miRNA targets rather than from nonspecific binding to unrelated transcripts. Together, these controls show that efficient miR-Space detection requires correct linker pairing and formation of the extended miRNA product.

Supplementary Fig. 2d extends this analysis to the 250-plex panel comprising 200 miRNAs and 50 mRNAs. Omitting the miRNA-extension step caused a marked reduction in miRNA reads, whereas mRNA detection was retained. This demonstrates that the no-extension control can be applied in a highly multiplexed panel and selectively assesses miRNA-detection specificity without compromising the mRNA component of the assay.

We next evaluated a direct 50-plex padlock-probe panel targeting native miRNAs without the extension strategy. As shown in Supplementary Fig. 2e, direct detection yielded few reads per cell and was strongly dominated by mmu-miR-124-3p. After excluding this highly abundant miRNA, the remaining targets yielded a mean of only 4.2 reads per cell, which was insufficient for robust single-cell analysis. Direct detection also failed to recover several spatial patterns observed with the extension-based assay. For example, Supplementary Fig. 2f shows that the extended mmu-miR-204-5p assay clearly detected enrichment along ventricular regions, whereas direct detection did not reproduce this distribution. Together with the more than twofold increase in extension-derived reads shown in main-text Fig. 2, these results demonstrate that miRNA extension substantially improves both detection efficiency and recovery of spatially informative miRNA patterns.

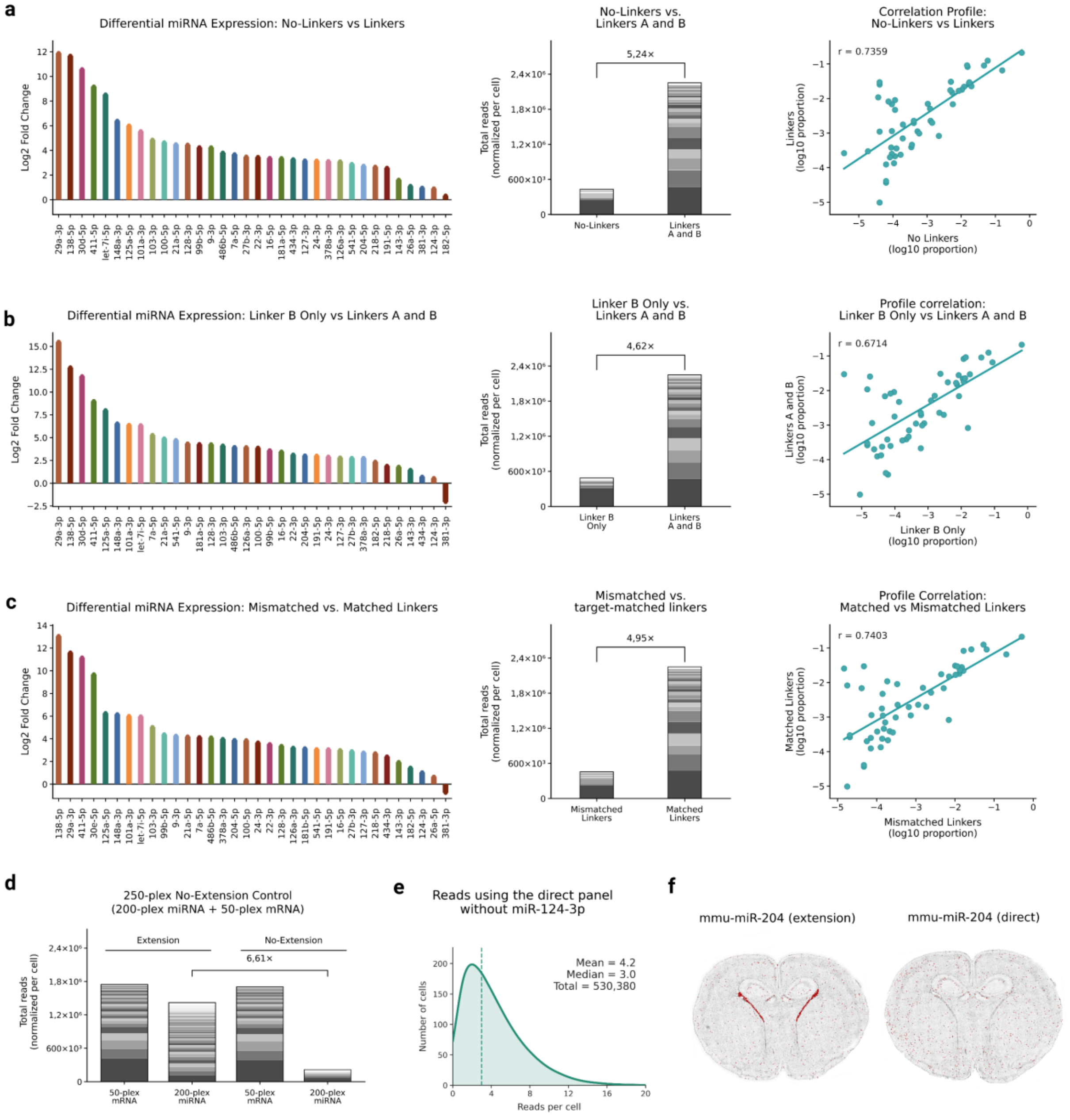


**Supporting Fig. 2 | No-extension and direct-detection controls confirm miR-Space specificity and sensitivity.**

a-c, The complete 50-plex extension reaction was compared with conditions lacking both linkers (a), containing linker B only (b), or using target-mismatched linker pairs (c). For each comparison, per-miRNA fold changes, total reads per cell, and correlations between relative abundance profiles are shown. Disruption of linker-mediated extension reduced total miRNA detection by approximately fivefold, although residual profiles remained positively correlated with the complete reaction. d, In the 250-plex panel comprising 200 miRNAs and 50 mRNAs, omission of miRNA extension markedly reduced miRNA reads while largely preserving mRNA detection. e, Direct padlock-probe detection without prior extension yielded a mean of 4.2 reads per cell after excluding mmu-miR-124-3p. f, Extension-based detection recovered ventricular enrichment of mmu-miR-204-5p, whereas direct detection did not reproduce this spatial pattern.

#### Supplementary Figure 3. miR-Space reproducibility and ligase compatibility

Supplementary Fig. 3a shows two additional miRNA expression profiles complementing Fig. 2a. These profiles were generated using panels targeting the same miRNAs but incorporating distinct barcode sets. Relative abundance was calculated as the proportion of the total signal contributed by each miRNA. Supplementary Fig. 3b extends the correlation analysis to all version 1 and version 2 datasets, as well as their mean profiles. The strong correlations observed across datasets show that different barcode identities yield similar miRNA expression profiles, supporting assay reproducibility across barcode sets. Across the seven datasets generated with versions 1 and 2, the mean number of reads per cell was 22.1 (Supplementary Fig. 3c). The observed variation likely reflects technical variability associated with tissue depth, experimental batch, and processing of sections obtained from three individual mice. We also evaluated alternative ligases for miRNA detection. SplintR successfully ligated linker B to mmu-miR-122-5p in HuH7 cells (Supplementary Fig. 3d). In mouse brain tissue, SplintR showed higher detection-probe ligation efficiency than KOD1Rnl (Supplementary Fig. 3e). Nevertheless, KOD1Rnl reproduced similar spatial patterns, indicating that the recovered miRNA distributions were robust to ligase choice.

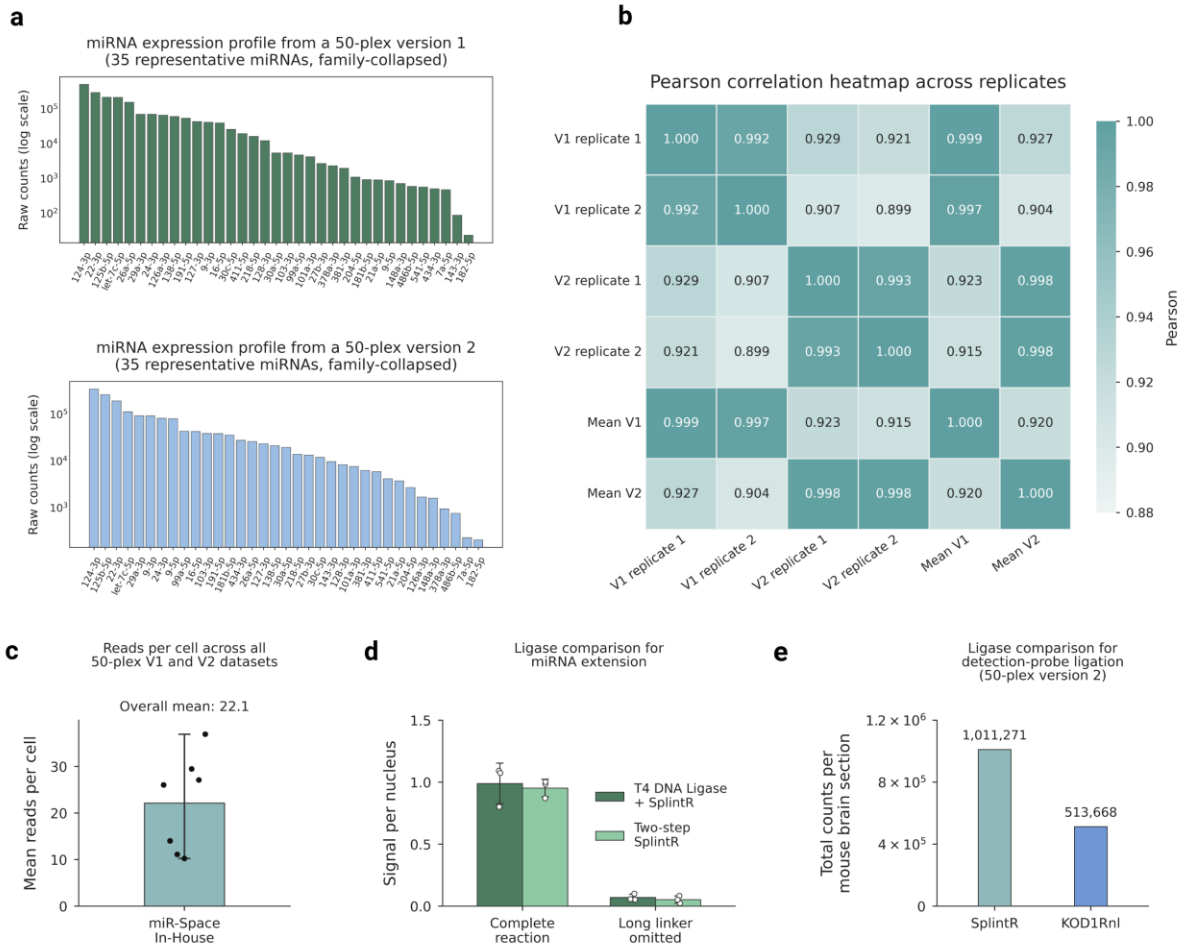


**Supplementary Fig. 3 | miR-Space reproducibility and ligase compatibility.**

a, Representative miRNA expression profiles obtained with the 50-plex version 1 and version 2 panels. Targets are ranked by abundance, and raw counts are shown on a logarithmic scale. b, Pearson correlation matrix comparing individual version 1 and version 2 replicates and their mean expression profiles. c, Mean reads per cell across seven 50-plex version 1 and version 2 datasets. The bar represents the mean across datasets, black dots indicate individual dataset means, and the error bar indicates the minimum-to-maximum range. d, Comparison of ligase configurations for miRNA extension. Bars show the mean signal per nucleus, error bars indicate standard deviation, and individual points represent replicates. Omission of the long linker served as a no-extension control. e, Comparison of SplintR and KOD1Rnl for detection-probe ligation in mouse brain sections. Bars show total detected counts per section.

#### Supplementary Figure 4. miR-Space USER version for enzymatic linker removal

Supplementary Fig. 4 presents an enzymatic strategy for removing the long linker after miRNA extension to expose the extended miRNA for subsequent padlock-probe hybridization. In the standard miR-Space workflow, the linker is removed by formamide washing. Here, we evaluated whether uracil-specific enzymatic cleavage could provide an alternative and whether pre-annealing linkers A and B before tissue application affected miRNA detection. As illustrated in Supplementary Fig. 4a, linker A first hybridizes to the endogenous miRNA, after which linker B hybridizes and is ligated to generate the extended miRNA product. The removable linker was designed to contain uracil residues, enabling selective cleavage after extension. Treatment with uracil-DNA glycosylase and Endonuclease III generated breaks at these positions, and subsequent formamide washing removed the resulting fragments, leaving the barcoded miRNA accessible for padlock-probe binding (Supplementary Fig. 4b). We refer to this approach as miR-Space USER, reflecting the use of uracil-specific excision for linker removal.

As indicated in main-text Fig. 2, we next compared four linker-treatment conditions: the standard workflow followed by formamide washing, pre-annealed linkers followed by formamide washing, the standard workflow combined with UDG and Endonuclease III treatment, and pre-annealed linkers combined with enzymatic cleavage. The reads-per-cell distributions include selected datasets also shown in Supplementary Fig. 3c, presented here to facilitate direct comparison of detection performance across experimental conditions. All four conditions produced broadly comparable signal distributions, with the pre-annealed USER condition yielding the highest number of reads per cell (Supplementary Fig. 4c). These results show that both sequential and pre-annealed linker delivery are compatible with miR-Space and that enzymatic cleavage preserves downstream miRNA detection while potentially improving signal recovery.

A targeted mmu-miR-122-5p experiment in HuH7 cells was then used to assess linker removal under different washing conditions (Supplementary Fig. 4d). UDG and Endonuclease III treatment preserved miRNA signal across the formamide concentrations tested. At 65% formamide, washing alone was insufficient for efficient linker removal, whereas USER-mediated cleavage maintained access to the extended miRNA target. These results indicate that the USER workflow can support milder formamide-washing conditions while preserving miRNA detection.

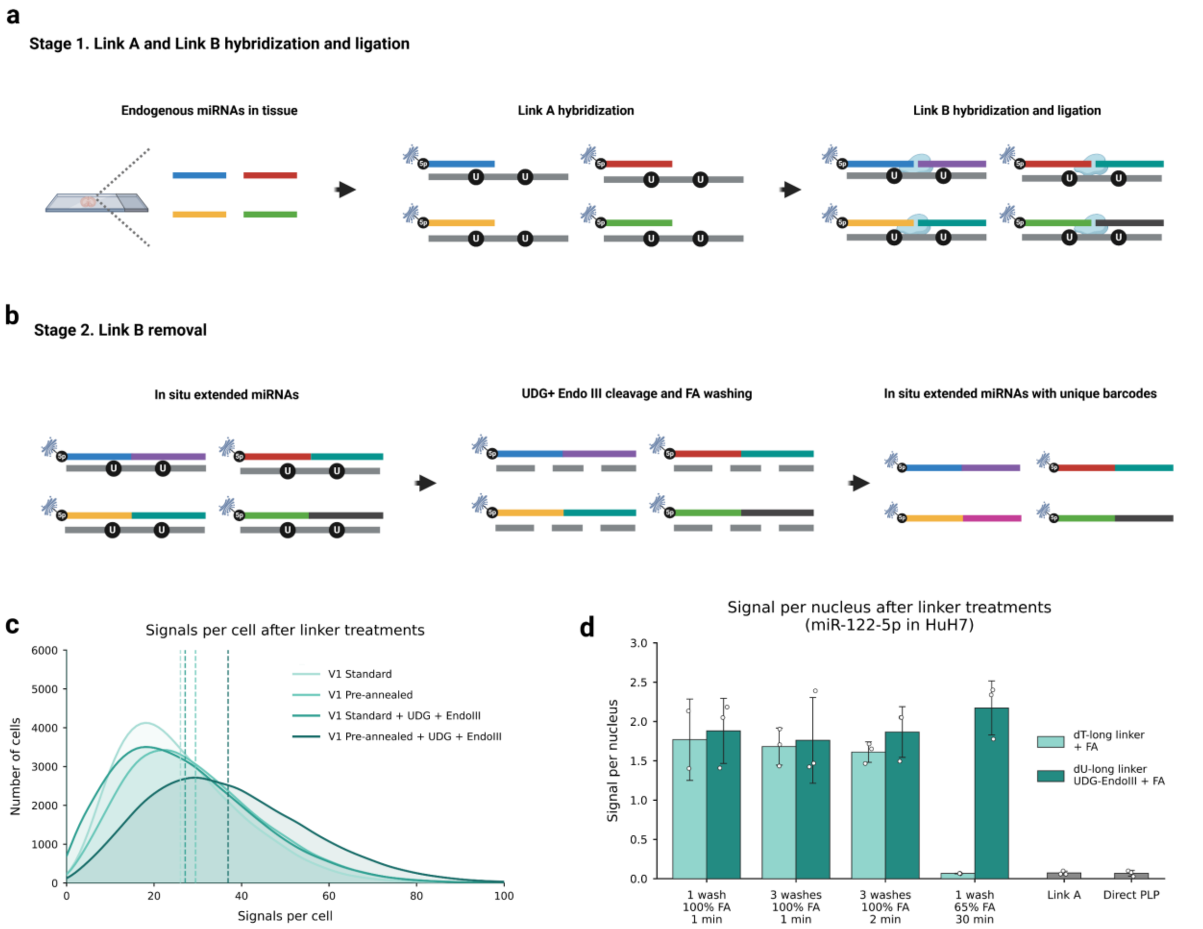


**Supporting Fig. 4 | miR-Space USER version for enzymatic linker removal.**

a, Schematic of miRNA extension. Linker A hybridizes to the endogenous miRNA, followed by linker B hybridization and ligation to generate the extended miRNA product. b, Linker B contains uracil residues that enable cleavage by uracil-DNA glycosylase and Endonuclease III. Subsequent washing removes the cleaved fragments and exposes the barcoded miRNA for padlock-probe hybridization. c, Reads-per-cell distributions obtained using sequential or pre-annealed linkers, with linker B removal by formamide washing alone or by enzymatic cleavage followed by washing. All conditions produced comparable signal distributions, with the pre-annealed and enzymatic cleavage yielding the highest signal. d, Detection of mmu-miR-122-5p in HuH7 cells after long-linker removal under different washing conditions. Enzymatic cleavage preserved signal across the tested formamide conditions and supported efficient linker removal under milder washing conditions.

#### Supplementary Figure 5. Crossplatform comparison of miR-Space Xenium with small RNA-seq

Supplementary Fig. 5 compares the miRNA expression profile obtained with the 50-plex miR-Space Xenium panel with three independent mouse brain small RNA-seq reference datasets: 4NBoost (Supplementary Fig. 5a), miRNATissueAtlas 2025 (Supplementary Fig. 5b), and GSE229981 (Supplementary Fig. 5c). For each reference, the abundance of each of the 50 targeted miRNAs was calculated relative to the total abundance of those same 50 miRNAs and log10-transformed. The left panels show positive correlations between miR-Space Xenium and all three references, with Pearson correlation coefficients of 0.76, 0.58, and 0.66, respectively. For GSE229981, three eligible biological replicates showed similar correlations, and the 8W-M1-Bulk sample is shown.

The middle panels plot the difference between platforms, calculated as miR-Space Xenium minus small RNA-seq relative abundance, against their mean relative abundance. Mean biases were negative for all comparisons (-0.38, -0.16, and -0.02), indicating slightly lower relative abundance in miR-Space Xenium, particularly compared with 4NBoost, whereas bias was nearly absent for GSE229981. The shallow fitted trends indicate no strong abundance-dependent disagreement. The right panels identify the most discordant miRNAs, including miR-7a-5p and members of the miR-34 family. These differences may reflect sample composition, as the reference datasets were generated from whole-brain bulk samples, whereas miR-Space Xenium measures spatially resolved abundance in individual tissue sections. miR-7a-5p and miR-34 family members are enriched in restricted hippocampal and ventricular regions, respectively, and may therefore contribute differently to overall abundance.

Supplementary Fig. 5d summarizes relative abundance across all four profiles, with miRNAs ordered by decreasing abundance in miR-Space Xenium. Overall, the positive correlations, small biases, and absence of strong abundance-dependent distortion support the ability of miR-Space Xenium to recover relative mature miRNA abundance across the targeted panel.

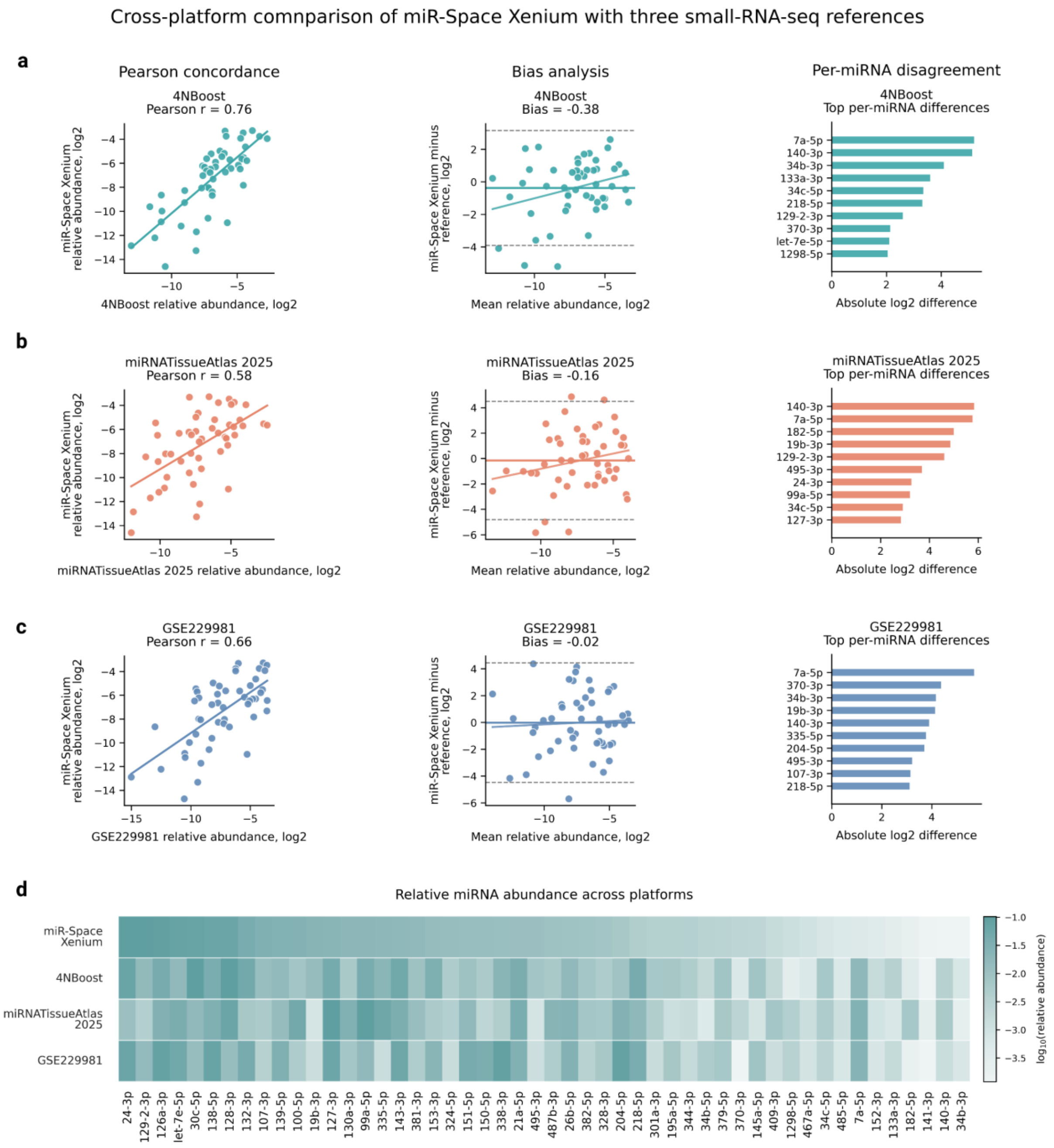


**Supplementary Figure 5 | Crossplatform comparison of miR-Space Xenium with small RNA-seq.** Relative mature miRNA abundance measured by miR-Space Xenium was compared with three independent mouse brain small RNA-seq reference datasets: 4NBoost (a), miRNATissueAtlas 2025 (b), and GSE229981 sample 8W-M1-Bulk (c). Within each dataset, abundance was normalized to the total signal across the 50 shared targets and log10-transformed. For each comparison, cross-platform agreement was assessed using Pearson correlation, bias analysis, and per-miRNA disagreement based on the absolute difference in log-transformed relative abundance. For GSE229981, eligible biological replicates were evaluated independently, and 8W-M1-Bulk was prespecified for display. The heatmap (d) shows log10-transformed relative abundance across the four profiles, with miRNAs ordered by decreasing abundance in miR-Space Xenium. Overall, miR-Space Xenium showed positive concordance with all three external references.

**Supplementary Figure 6. Spatial atlas of individual miRNA expression across the mouse brain**

Supplementary Fig. 6 presents coronal and sagittal sections illustrating distinct regional patterns of mature miRNA expression. miR-24-3p and miR-128-3p are preferentially enriched in the cerebral cortex. miR-138-5p and miR-218-5p show increased abundance in the caudoputamen/striatum and hippocampus, whereas miR-139-5p and miR-153-3p are preferentially enriched in the striatum. miR-338-3p is strongly associated with major fiber tracts, while miR-21a-5p shows a broader distribution with relative enrichment in fiber-tract regions.

miR-126a-3p displays a characteristic endothelial pattern, delineating elongated vessel-like structures. miR-150-5p shows a related, although not completely overlapping, vascular-associated distribution, whereas miR-143-3p is prominently enriched in the vasculature. miR-204-5p, miR-1298-5p, and miR-34c-5p show ventricular-associated expression. miR-204-5p and miR-1298-5p predominantly label the lateral and third ventricles, with little or no signal in the fourth ventricle, whereas miR-34c-5p labels the complete ventricular system, including the fourth ventricle.

miR-495-3p and miR-382-5p are preferentially enriched in ventral regions of the coronal sections. As shown in main-text Fig. 3, miR-195a-5p prominently marks the cerebellum and also shows enrichment in the dentate gyrus, a pattern likewise observed in the coronal section. miR-182-5p is enriched in piriform cortical areas, whereas miR-133a-3p shows a spatially restricted pattern in the reticular nucleus of the thalamus.

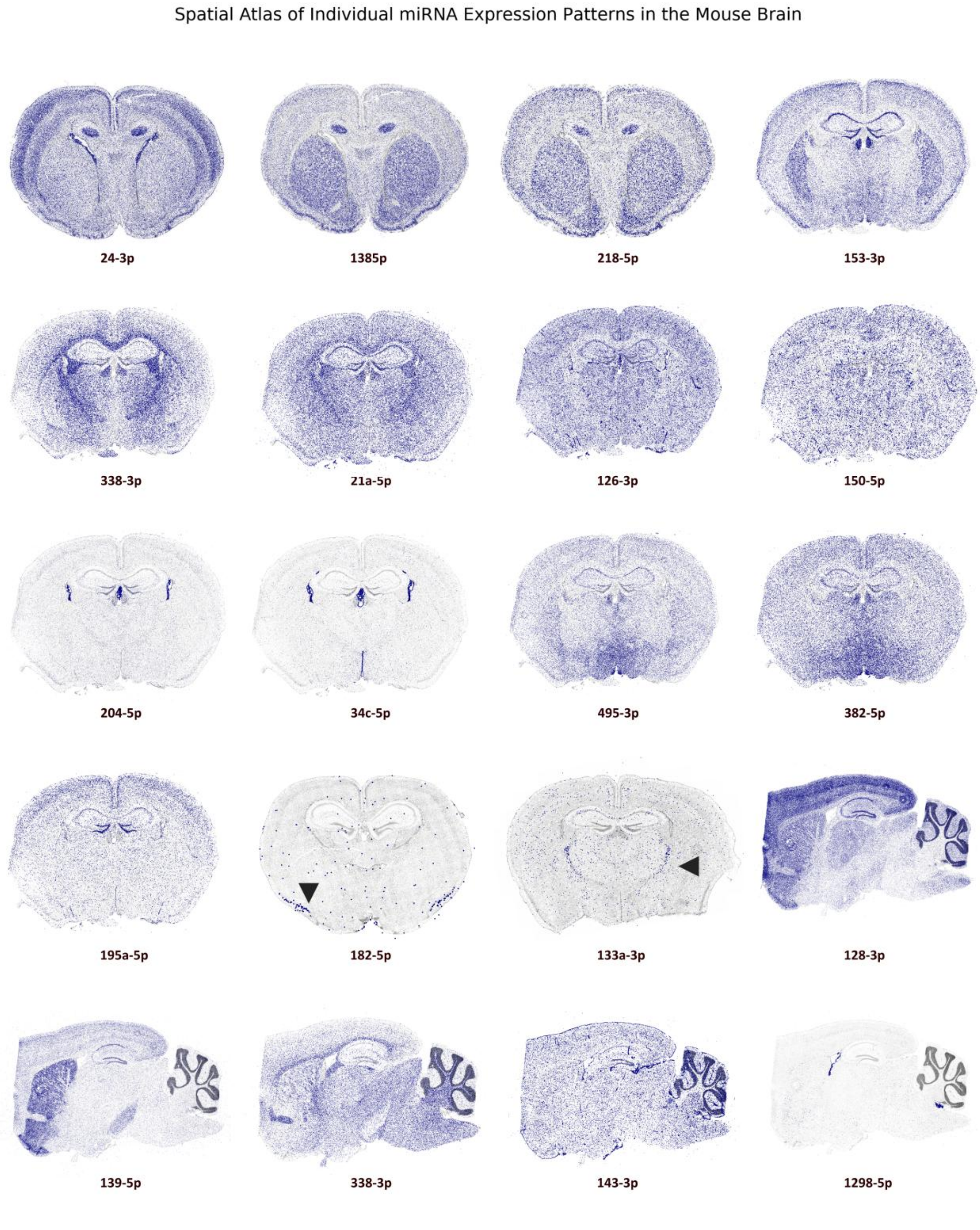


**Supplementary Fig. 6. Spatial atlas of individual miRNA expression across the mouse brain.** Representative miRNAs exhibit distinct regional expression patterns, with enrichment across the cortex, striatum, fiber tracts, vasculature, ventricular system, ventral brain regions, dentate gyrus, piriform cortex, and thalamic reticular nucleus.

**Supplementary Figure 7. Single-cell analysis of mRNA and miRNA expression and candidate miRNA-mRNA relationships**

Supplementary Fig. 7 presents an integrated characterization of cell identities, molecular complexity, assay specificity, and joint mRNA–miRNA expression obtained using miR-Space. Supplementary Fig. 7a shows cell-type annotation based on the expression of 247 mRNA marker genes, resolving major neuronal, glial, vascular, oligodendroglial, and ventricular-associated populations. Supplementary Fig. 7b summarizes total mRNA counts and the number of detected mRNA features per cell, providing measures of transcript recovery and molecular complexity across annotated populations. Supplementary Fig. 7c shows that clustering based exclusively on the 50-plex miRNA panel separates major brain cell classes, indicating that multiplexed miRNA profiles contain sufficient information to distinguish broad cellular identities. Supplementary Fig. 7d compares total miRNA counts and detected miRNA features across cell types. Supplementary Fig. 7e shows that the in-house no-extension control, generated using the same miRNA panel as the miR-Space Xenium experiment, yielded markedly fewer normalized reads than the positive control, supporting the dependence of miR-Space detection on miRNA extension. Supplementary Fig. 7f compares cell-type composition between two consecutive tissue sections from the same mouse, showing broadly consistent representation of major populations.

Supplementary Fig. 7g explores spatial relationships between miRNAs and reported mRNA targets. miR-495-3p was enriched in diencephalic populations, whereas Arc mRNA was more abundant in cortical and hippocampal excitatory neurons. Similarly, miR-204-5p was enriched in choroid plexus-associated cells, whereas Meis2 was more abundant in medium spiny neurons. Arc and Meis2 have been reported as direct targets of miR-495 and miR-204, respectively (Bastle et al., 2018; Conte et al., 2010). These reciprocal spatial patterns are consistent with previously reported regulatory relationships and illustrate how joint miRNA-mRNA profiling with miR-Space can be used to explore candidate post-transcriptional interactions across anatomically defined cell populations.

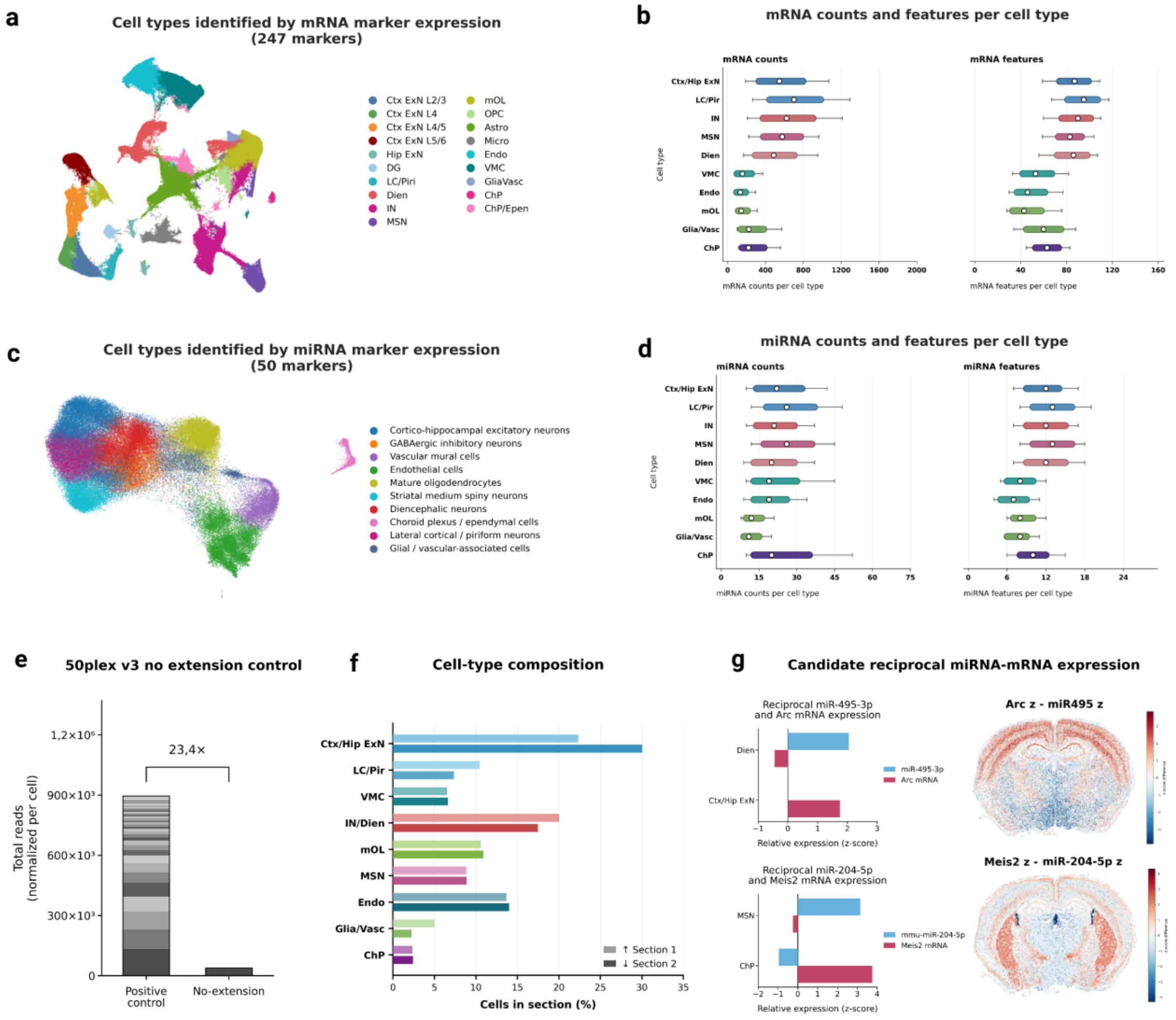


**Supplementary Fig. 7 | Cell type characterization of mRNA and miRNA expression and candidate miRNA-mRNA relationships.**

a, Cell-type annotation based on the expression of 247 mRNA marker genes. b, Distribution of total mRNA counts and detected mRNA features per cell across annotated cell types. c, Cell-type annotation based exclusively on the expression of 50 miRNAs. d, Distribution of total miRNA counts and detected miRNA features per cell across cell types. e, Comparison of normalized total reads between the positive control and an in-house no-extension control generated using the same 50-plex v3 panel. f, Relative abundance of identified cell types in the two analyzed tissue sections. g, Reciprocal expression and spatial distributions of miR-495-3p and Arc, and miR-204-5p and Meis2, across cell populations.

#### Supplementary Figure 8. Spatial distribution of miRNA-defined cell types across the mouse brain

Supplementary Fig. 8 shows the spatial distribution of the 12 cell populations identified from miRNA expression across the mouse brain section. After defining miRNA-associated cell populations using Wilcoxon differential-expression analysis, their projection onto the tissue revealed distinct anatomical enrichment patterns. These included cortical enrichment of excitatory neuronal populations, striatal enrichment of medium spiny neurons, ventral enrichment of diencephalic populations, and restricted ventricular localization of choroid plexus and ependymal cells. Inhibitory, endothelial, mural, oligodendroglial, glial-associated, lateral cortical/piriform, cortico-hippocampal, and thalamic/subcortical populations also showed characteristic regional distributions. As defined in Supplementary Tables 3 and 4, the miRNA-associated cell populations were subsequently classified as either higher-confidence populations or mixed/broader populations. Supplementary Fig. 8 displays their spatial distributions according to this classification. Together, these maps show that miRNA-based cell-type annotation recapitulates the principal anatomical organization of major neuronal and non-neuronal populations.

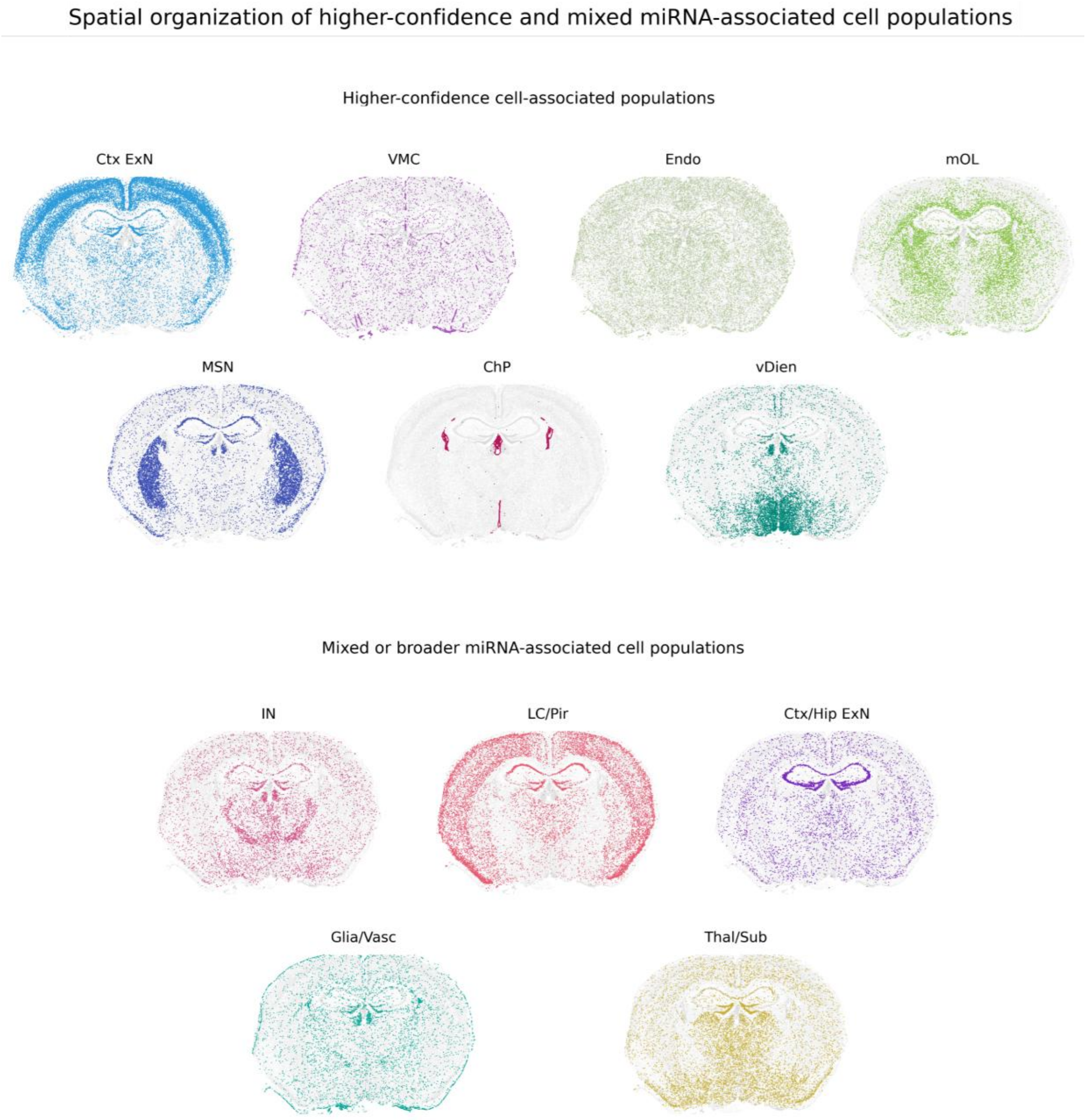


**Supplementary Fig. 8 | Principal spatial enrichment of miRNA-defined cell types across the mouse brain.** Individual spatial masks show the main anatomical enrichment patterns of 12 cell populations identified from miRNA expression, including cortical and cortico-hippocampal excitatory neurons, inhibitory neurons, vascular mural and endothelial cells, mature oligodendrocytes, striatal medium spiny neurons, ventral diencephalic neurons, choroid plexus and ependymal cells, lateral cortical and piriform neurons, glial and vascular-associated cells, and thalamic and subcortical populations.

**Supplementary Figure 9. Reproducibility and cell-type specificity of high-confidence miRNA signatures across independent datasets**

To assess the robustness of the cell-associated miRNA signatures across independent experiments, Supplementary Figure 9 compares the six high-confidence modules across the six spatial miRNA datasets. For each dataset, mean module scores were calculated across the six corresponding cell populations: cortical excitatory neurons, vascular/endothelial cells, mature oligodendrocytes, striatal medium spiny neurons, choroid plexus/ependymal cells, and ventral diencephalic neurons. Despite differences in developmental stage, experimental replicate, and multiplexed workflows, each module consistently reached its highest or preferential score in the expected cellular population. The cortical excitatory, vascular, oligodendrocyte, striatal MSN, choroid plexus, and ventral diencephalic signatures retained their characteristic population-specific profiles across datasets, while non-target populations generally showed substantially lower module scores. These results demonstrate that the identified miRNA signatures are reproducible across biological samples and experimental configurations and support their robustness as cell-associated molecular features of the corresponding brain populations.

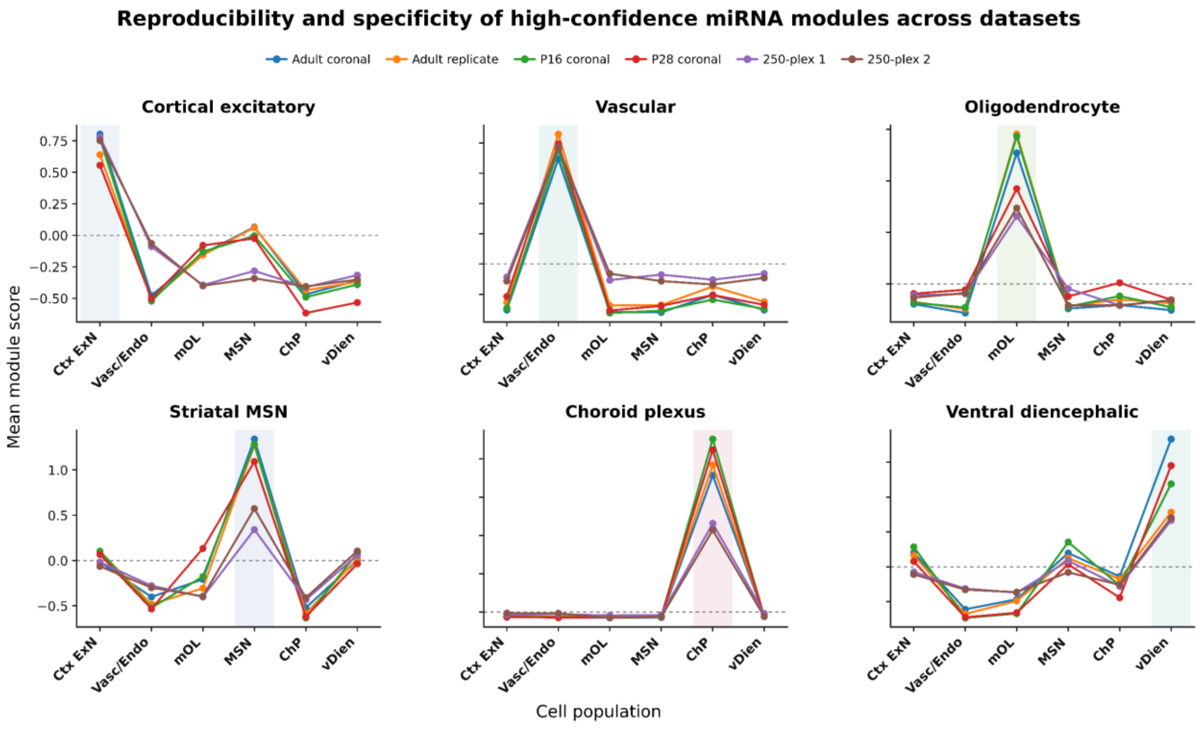


**Supplementary Figure 9 | Reproducibility and cell-type specificity of high-confidence miRNA signatures across independent datasets.**

Mean module scores for the six high-confidence miRNA signatures are shown across six corresponding cell populations in six independent spatial miRNA datasets: Adult coronal, Adult replicate, P16 coronal, P28 coronal, 250-plex 1, and 250-plex 2. Each panel represents one miRNA signature, and colored lines indicate individual datasets. Cell populations are ordered as cortical excitatory neurons (Ctx ExN), vascular/endothelial cells (Vasc/Endo), mature oligodendrocytes (mOL), striatal medium spiny neurons (MSN), choroid plexus/ependymal cells (ChP), and ventral diencephalic neurons (vDien). Shaded regions indicate the expected target population for each signature. Across datasets, the cortical excitatory, vascular, oligodendrocyte, striatal MSN, choroid plexus, and ventral diencephalic signatures showed preferential enrichment in their corresponding target populations, supporting the reproducibility and specificity of these cell-associated miRNA programs across biological samples and profiling workflows.

#### Supplementary Figure 10. ell-type specificity of high-confidence miRNA modules across annotated adult brain cell types

To further evaluate the cellular specificity of the six high-confidence miRNA modules, Supplementary Fig. 10 compares per-cell module score distributions across all twelve annotated cell populations in the representative adult coronal dataset. Each module showed preferential enrichment in its expected target population, while scores were substantially lower across most non-target cell types. The cortical excitatory, vascular/endothelial, mature oligodendrocyte, striatal MSN, and choroid plexus/ependymal modules showed particularly strong discrimination of their corresponding cellular populations, with ROC AUC values ranging from 0.90 to 1.00. For all six modules, target-cell scores were significantly higher than those observed across pooled non-target populations after Benjamini–Hochberg correction. Together, these results support the cell-type specificity of the identified miRNA modules and show that their enrichment is maintained when evaluated against the broader diversity of annotated adult brain cell populations.

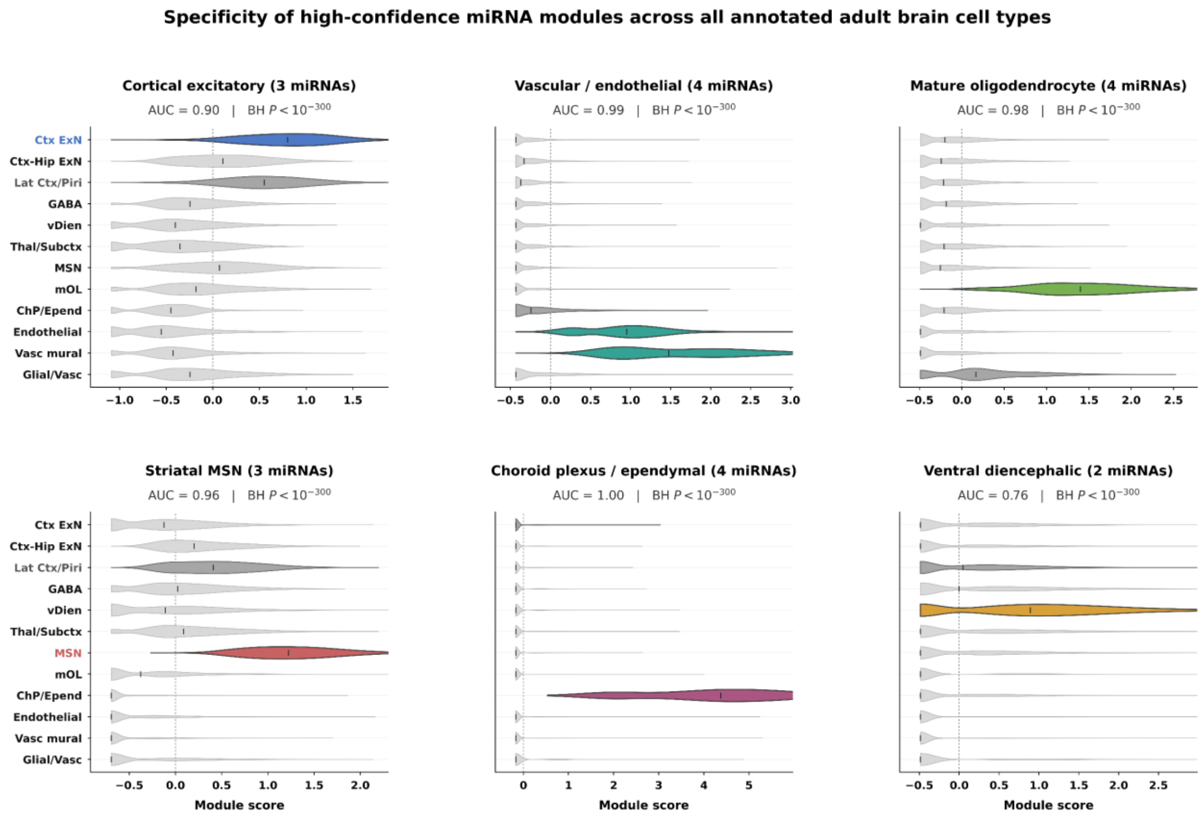


**Supplementary Figure 10 | Cell-type specificity of high-confidence miRNA modules.** Per-cell module scores are shown for six high-confidence miRNA modules across twelve annotated adult brain cell populations. Colored violins indicate the expected target population(s), and grey violins indicate non-target populations; for the vascular/endothelial module, both endothelial and vascular mural cells were considered targets. Black lines indicate median scores. Specificity was assessed using a one-sided Mann–Whitney U test comparing target versus pooled non-target cells, with Benjamini–Hochberg correction across modules. ROC AUC values indicate target-versus-non-target discrimination. All six modules were preferentially enriched in their expected cell populations.

#### Supplementary Table 4. Comparison of miR-Space with spatial transcriptomics technologies

Supplementary Table 4 positions miR-Space within the current landscape of spatial transcriptomic technologies capable of measuring miRNAs directly in intact tissue. These approaches differ substantially in molecular coverage, spatial resolution, tissue compatibility, and support for cell-level interpretation. Sequencing-based platforms such as STRS and Patho-DBiT provide broad transcriptome-wide profiling, including non-coding RNAs, but rely on spot- or pixel-level measurements and require computational deconvolution or reconstruction for cellular interpretation. Spectrum-FISH enables multiplexed, sequencing-free imaging of selected miRNAs, although it has been demonstrated with a smaller panel and in a more restricted tissue context.

Within this landscape, miR-Space combines substantially higher miRNA multiplexing with direct single-cell assignment across multiple cell types while preserving spatial organization in intact tissue. Supplementary Table 4 was restricted to platforms that generate multiplexed, spatially indexed RNA-expression profiles directly from intact tissue. Conventional LNA-ISH, miRNAscope, targeted nanowell assays, and miRNA FISH methods limited to individual miRNAs or small miRNA sets were excluded because they are primarily designed for targeted localization or validation rather than broad spatial transcriptomic profiling. Supplementary Table 4 summarizes the principal characteristics, capabilities, and limitations of the selected approaches.

| Comparison criterion | STRS | Patho-DBiT | Spectrum-FISH | miR-Space |
|---|---|---|---|---|
| **RNA coverage** | Transcriptome-wide total RNA, including non-coding and non-polyadenylated RNAs. | Transcriptome-wide total RNA, including non-coding and non-polyadenylated RNAs. | Targeted panel: 24 miRNAs and 9 m6A-modified mRNAs. | Targeted panel; 200 miRNAs and 50 mRNAs demonstrated. |
| **Spatial resolution** | 55 µm Visium spots. | 10-50 µm pixels. | Single-molecule fluorescence imaging. | Single-molecule and single-cell resolution. |
| **Single-cell resolution** | No; spot-level measurements. | No; pixel-level measurements with computational cell assignment. | Limited; cell-level measurements demonstrated in selected NeuN-positive cells. | Yes; cell-level assignment across multiple cell types |
| **Tissue** | Fresh-frozen. | FFPE archival tissue. | Acute olfactory-bulb slices. | Fresh-frozen mouse and human brain. |
| **Commercialization** | Adaptable to Visium. | No commercial kit; custom microfluidics. | No commercial kit; custom device. | Adaptable to Xenium. |
| **Main capability** | Broad spatial profiling beyond poly(A)-based capture. | Broad spatial profiling in FFPE tissue. | Multiplexed, sequencing-free imaging of selected targets. | Highly multiplexed miRNA detection with no-extension controls and cellular context. |
| **Main constraint** | Limited cellular resolution and mature-miRNA sensitivity. | Complex workflow and computational reconstruction. | Lower demonstrated plex and validation limited to a olfactory-bulb region. | Targeted design requiring custom probes and panel-specific validation. |
| **Reference** | McKellar et al. (2023), Nature Biotechnology. | Bai et al. (2024), Cell. | Ji et al. (2026), Nature Biomedical Engineering. | Robles-Remacho et al., this study. |

**Supplementary Table 4. Comparison of miR-Space with spatial transcriptomics technologies.** STRS and Patho-DBiT provide broad transcriptome-wide RNA coverage using spatially indexed sequencing, whereas Spectrum-FISH and miR-Space use targeted approaches for direct spatial detection of selected miRNAs. The platforms are compared according to RNA coverage, spatial resolution, cell-level analysis, tissue compatibility, commercialization, principal capability, main limitation, and reference.

**Supplementary Table 5. Literature-based validation and novelty of miR-Space spatial miRNA patterns**

Supplementary Table 5 provides a systematic literature-based assessment of the spatial miRNA expression patterns identified by miR-Space. Patterns observed with the 50-plex miRNA panel implemented on the Xenium platform were compared with previously published evidence and assigned to one of three standardized categories: Confirmed, Partial regional support, or Candidate novel. Confirmed indicates direct agreement with previously reported anatomical or cell-type-specific localization. Partial regional support indicates that previous studies support expression in the brain, a related anatomical region, developmental context, or cell type, but do not reproduce the complete spatial pattern observed here. Candidate novel identifies spatial distributions for which no directly matching localization was found in the reviewed literature. These candidate novel patterns should therefore be interpreted as potentially new spatial associations revealed by the miR-Space Xenium 50-plex analysis, rather than as evidence that the corresponding miRNAs have never previously been detected in brain tissue.

| miRNA | miR-Space localization | Supporting study / interpretation |
|---|---|---|
| **miR-24-3p** | Broad neuronal expression, with cortical enrichment | Neuronal and cortical expression has been reported, but specific cortical enrichment remains unconfirmed (Kang et al., Cell Mol Life Sci, 2019; Higuchi et al., Nat Commun, 2025). |
| **miR-487b-3p** | Broad expression, with ventral enrichment | Broad mammalian and brain expression has been reported, but specific ventral enrichment remains unconfirmed (Landgraf et al., Cell, 2007; Engel et al., Nat Commun, 2025). |
| **miR-129-2-3p** | Broad neuronal expression, with reduced signal in fiber tracts | miR-129-3p expression in the developing neocortex and cortical plate has been reported, supporting a role in neuronal migration (Wu et al., Cell Death Dis, 2019). |
| **miR-26b-5p** | Broad neuronal expression | Abundant mouse-brain expression is documented, although an adult cell-resolved map is lacking (Landgraf et al., Cell, 2007). |
| **miR-126a-3p** | Endothelial and vascular enrichment | Endothelial-restricted expression and vascular functions were demonstrated in mice (Wang et al., Dev Cell, 2008). |
| **miR-382-5p** | Broad expression, with ventral enrichment | Brain-enriched expression and detection in human olfactory neuronal tissue have been reported, but specific ventral brain enrichment remains unconfirmed (Mor et al., Neurobiol Dis, 2013). |
| **let-7e-5p** | Broad neuronal expression | Broad expression across the adult mouse CNS has been reported (Bak et al., RNA, 2008). |
| **miR-328-3p** | Ventral brain enrichment | Brain expression is present in mammalian miRNA atlases, but a ventrally enriched pattern has not been described (Landgraf et al., Cell, 2007). |
| **miR-30c-5p** | Broad neuronal expression | miR-30c expression in the adult SVZ, olfactory bulb, and hippocampus has been reported (Sun et al., Cell Prolif, 2016). |
| **miR-204-5p** | Choroid plexus and ventricular enrichment | The adult lateral-ventricle choroid plexus was identified as a major source of miR-204 (Lepko et al., EMBO J, 2019). |
| **miR-138-5p** | Broad neuronal expression, with hippocampal and striatal enrichment | Neuronal expression in the neocortex, hippocampus, and cerebellum has been reported, but specific striatal enrichment remains unconfirmed (Obernosterer et al., RNA, 2006). |
| **miR-218-5p** | Broad neuronal expression, with hippocampal and striatal enrichment | High expression in the hippocampus and enrichment in excitatory principal neurons and GABAergic interneurons have been demonstrated; specific striatal enrichment remains unconfirmed (Taylor et al., eLife, 2023). |
| **miR-128-3p** | Cortical enrichment | Brain-enriched expression and prominent localization in cortical neurons have been demonstrated (Tan et al., Science, 2013; Franzoni et al., eLife, 2015). |
| **miR-301a-3p** | Broad neuronal expression | miR-301a-3p was experimentally detected in a mammalian small-RNA expression atlas, but broad neuronal and cell-resolved brain expression remain unconfirmed (Landgraf et al., Cell, 2007). |
| **miR-132-3p** | Cortical enrichment | miR-132 was directly studied in visual cortex and included in cell-type-resolved brain miRNA profiling, but cortex-specific enrichment remains unconfirmed (Mellios et al., Nat Neurosci, 2011; He et al., Neuron, 2012). |
| **miR-195a-5p** | Cerebellar enrichment | Adult mouse CNS profiling showed strong cerebellar enrichment (Bak et al., RNA, 2008). |
| **miR-107-3p** | Broad neuronal expression. | miR-107 expression in rodent hippocampus and hippocampal neurons has been demonstrated, but broad brain-wide neuronal expression remains unconfirmed (Redell et al., J Neurosci Res, 2009; Wang et al., Am J Pathol, 2010). |
| **miR-344-3p** | Broad neuronal expression, with striatal enrichment | miR-344-3p expression has been demonstrated in the developing CNS and selected adult brain regions, but adult striatal enrichment remains unconfirmed (Liu et al., J Mol Histol, 2014). |
| **miR-139-5p** | Striatal and medium spiny neuron enrichment | Hippocampal expression and functional effects on neurogenesis have been reported, but enrichment in adult striatum or medium spiny neurons remains unconfirmed (Liu et al., Transl Psychiatry, 2021). |
| **miR-34b-5p** | Choroid plexus and ventricular enrichment | miR-34b-5p has been detected in rodent hippocampus and hippocampal astrocytes, but adult choroid-plexus or ventricular enrichment remains unconfirmed (Liu et al., BMC Neurosci, 2016). |
| **miR-100-5p** | White-matter and mature oligodendrocyte enrichment | miR-100 was included in oligodendrocyte-lineage profiling, but enrichment in mature oligodendrocytes or white matter remains unconfirmed (Lau et al., J Neurosci, 2008). |

| miRNA | miR-Space localization | Supporting study / interpretation |
|---|---|---|
| **miR-379-5p** | Broad neuronal expression | miR-379-5p belongs to the brain-enriched, neuronally regulated miR-379/410 cluster, but its broad neuronal distribution has not been independently established (Fiore et al., EMBO J, 2009). |
| **miR-19b-3p** | Broad neuronal expression | miR-19b-3p has been detected in mammalian neuronal libraries and cell-type-resolved brain profiles, but broad adult neuronal distribution remains unconfirmed (Landgraf et al., Cell, 2007; He et al., Neuron, 2012). |
| **miR-370-3p** | Broad neuronal expression | miR-370-3p expression has been detected in the developing mouse brain, but broad neuronal or adult brain-wide expression remains unconfirmed (Liu et al., FEBS Lett, 2013). |
| **miR-127-3p** | Broad neuronal expression, with ventral enrichment | Neuronal expression and functions in neurite growth and neuronal survival have been reported, but enrichment in ventral brain regions remains unconfirmed (He et al., Neuron, 2012; Zhou et al., Sci Rep, 2016). |
| **miR-145a-5p** | Vascular mural-cell enrichment | miR-145a-5p is strongly associated with vascular smooth-muscle cells and pericytes and regulates smooth-muscle phenotype (Xin et al., *Genes Dev*, 2009; Larsson et al., Genome Med, 2009). |
| **miR-130a-3p** | Broad neuronal expression, with ventral enrichment | miR-130a expression in developing cortical neurons and its regulation of neurite and dendritic-spine development have been demonstrated, but broad adult neuronal or ventral brain enrichment remains unconfirmed (Zhang et al., Protein Cell, 2016). |
| **miR-409-5p** | Broad neuronal expression | miR-409-5p has been detected in mouse cortex and primary hippocampal neurons, but broad neuronal or brain-wide spatial expression remains unconfirmed (Guo et al., Front Neurosci, 2019). |
| **miR-99a-5p** | Broad neuronal expression | miR-99a-5p has been detected in the adult hippocampus and functionally studied in neurons after cerebral ischemia, but broad neuronal and cell-resolved brain expression remain unconfirmed (Eacker et al., PLoS One, 2011; Tao et al., J Neurol Sci, 2015). |
| **miR-1298-5p** | Choroid plexus and ventricular enrichment | miR-1298-5p has been detected in human neocortical and neurological-disease samples, but adult choroid-plexus or ventricular enrichment remains unconfirmed (Baloun et al., Epilepsy Res, 2020). |
| **miR-335-5p** | Broad neuronal expression, with ventral enrichment | miR-335-5p is brain enriched, expressed in excitatory and inhibitory neurons, and regulates neuronal excitability; specific ventral or hypothalamic enrichment remains unconfirmed (Heiland et al., Proc Natl Acad Sci USA, 2023). |
| **miR-467a-5p** | Broad neuronal expression | miR-467a-5p has been identified in mouse small-RNA sequencing datasets, but physiological brain expression and broad adult neuronal localization remain unconfirmed (Chiang et al., Genes Dev, 2010). |
| **miR-143-3p** | Vascular mural-cell enrichment | miR-143-3p is strongly associated with vascular smooth-muscle cells, consistent with mural-cell enrichment (Xin et al., Genes Dev, 2009). |
| **miR-34c-5p** | Ventricular-system enrichment | miR-34c is represented in neural miRNA profiles, but normal adult ventricular enrichment has not been reported (Landgraf et al., Cell, 2007). |
| **miR-381-3p** | Broad expression, with ventral enrichment | miR-381-3p belongs to the brain-enriched and neuronally regulated miR-379/410 cluster, but its broad neuronal distribution and ventral enrichment remain unconfirmed (Fiore et al., EMBO J, 2009). |
| **miR-485-5p** | Broad neuronal expression | miR-485-5p has been detected in hippocampal and cortical neurons, but broad neuronal or brain-wide expression remains unconfirmed (He et al., Transl Neurosci, 2021; Cheng et al., Mol Neurobiol, 2021). |
| **miR-153-3p** | Striatal and medium spiny neuron enrichment | miR-153-3p regulates neural stem-cell proliferation and differentiation, but enrichment in adult striatum remains unconfirmed (Qiao et al., Cell Death Differ, 2020; Dong et al., *Aging*, 2023). |
| **miR-7a-5p** | Hypothalamic enrichment | Hypothalamic enrichment of miR-7a has been demonstrated, with prominent expression in several hypothalamic nuclei (Herzer et al., J Neuroendocrinol, 2012). |
| **miR-324-5p** | Broad neuronal expression, with striatal enrichment | Hippocampal association and a functional role in spatial-memory formation have been demonstrated, but broad neuronal or striatal enrichment remains unconfirmed (Busto et al., Mol Neurobiol, 2016). |
| **miR-152-3p** | Meningeal surface enrichment | miR-152-3p has been detected in cerebrospinal-fluid exosomes and studied in vascular smooth-muscle cells, but meningeal enrichment remains unconfirmed (Li et al., Folia Neuropathol, 2022). |

| miRNA | miR-Space localization | Supporting study / interpretation |
|---|---|---|
| **miR-151-5p** | Broad neuronal expression | miR-151-5p has been detected in mouse cerebrovasculature and developing cortical neural stem cells (Chum et al., J Alzheimers Dis, 2022; Peng et al., Stem Cell Reports, 2026). |
| **miR-133a-3p** | Inhibitory neuron and internal-capsule enrichment | miR-133a-3p is primarily recognized as a muscle-enriched miRNA and has been examined in mouse cerebrovascular profiles, but inhibitory-neuron or internal-capsule enrichment remains unconfirmed (Chum et al., J Alzheimers Dis, 2022). |
| **miR-150-5p** | Endothelial and vascular enrichment | miR-150-5p has been detected and functionally studied in brain microvascular endothelial cells, but endothelial or vascular enrichment remains unconfirmed (Perales et al., Alcohol Clin Exp Res, 2022). |
| **miR-182-5p** | Piriform neuron enrichment | miR-182-5p has been detected in olfactory tissues and described as neuron enriched, but adult piriform-cortex enrichment remains unconfirmed (Choi et al., Neuron, 2008; Wang et al., Front Cell Neurosci, 2017). |
| **miR-338-3p** | White-matter and mature oligodendrocyte enrichment | miR-338 is enriched in oligodendrocytes and promotes oligodendrocyte differentiation (Lau et al., J Neurosci, 2008; Zhao et al., Neuron, 2010). |
| **miR-141-3p** | Olfactory-bulb enrichment | The miR-200 family, including miR-141, regulates olfactory neurogenesis and is enriched in olfactory and pituitary tissues (Choi et al., Neuron, 2008; Landgraf et al., Cell, 2007). |
| **miR-21a-5p** | Broad expression, with white-matter and fiber-tract enrichment | miR-21a-5p is abundantly expressed in oligodendrocytes and reactive neural cells, but normal adult white-matter or fiber-tract enrichment remains unconfirmed (Pathania et al., Psychiatry Res, 2017). |
| **miR-140-3p** | Broad neuronal expression | miR-140-3p has been detected in embryonic cortical neurons, but broad adult neuronal and brain-wide localization remain unconfirmed (Kim et al., Proc Natl Acad Sci USA, 2004). |
| **miR-495-3p** | Ventral diencephalic neuron enrichment | miR-495-3p belongs to the brain-enriched miR-379/410 cluster and is expressed in the developing cerebral cortex, but ventral diencephalic neuronal enrichment remains unconfirmed (Fiore et al., EMBO J, 2009; Wang et al., Int J Biol Sci, 2024). |
| **miR-34b-3p** | Choroid plexus and ventricular enrichment | miR-34b-3p has been detected in the adult mouse hippocampus, but normal choroid-plexus or ventricular-system enrichment remains unconfirmed (Zhang et al., Neurosci Bull, 2023). |

**Supporting Table 5. Literature-based validation and novelty of miR-Space spatial miRNA patterns.** Reported expression patterns were evaluated against primary studies and classified according to the strength of available evidence. “Confirmed” indicates direct regional or cell-type support, “partial regional support” indicates indirect or incomplete evidence, and “not previously reported” indicates that the proposed localization has not been demonstrated.

#### Supplementary Table 6. Integrated interpretation of miRNA-defined cell populations and associated miRNA signatures

Supplementary Table 6 integrates mRNA-based cell identity, quantitative miRNA enrichment, and spatial concordance across the 12 annotated cell populations. For each population, the table reports the number of cells, principal supporting mRNA markers, the most enriched miRNAs with their corresponding scores and log2 fold changes, and an integrated interpretation of the molecular and anatomical evidence. High-confidence populations are specifically indicated and include endothelial cells, cortical excitatory neurons, ventral diencephalic neurons, striatal medium spiny neurons, mature oligodendrocytes, choroid plexus/ependymal cells, and vascular mural cells. In contrast, cortico-hippocampal, lateral cortical/piriform, thalamic/subcortical, and glial/vascular-associated populations showed broader or less specific miRNA profiles, with their annotation supported primarily by mRNA expression and spatial localization.

| Cell population (n cells) | mRNA support | Primary miRNA marker(s) [score; log2FC] | Integrated conclusion |
|---|---|---|---|
| **Endothelial cells (n = 16,143)** | Strong endothelial support, including Cldn5, Pecam1, Emcn, Kdr, Acvrl1, Adgrl4, Cd93, Sox17, Fgd5 and Nostrin. | miR-126a-3p [151.4; 3.87]. miR-150-5p [117.2; 3.83] showed vascular-associated enrichment but lower endothelial specificity. | High-confidence endothelial population supported by concordant mRNA and miRNA expression. miR-126a-3p represented the principal endothelial-associated miRNA, whereas miR-150-5p provided additional vascular support. |
| **Cortical excitatory neurons (n = 20,714)** | Strong cortical excitatory support, including Satb2, Slc17a7, Neurod6, Nrn1, Arc and Epha4. Lamp5 expression indicated a degree of cellular heterogeneity. | miR-128-3p [123.2; 2.04] and miR-132-3p [86.1; 1.74]. miR-24-3p [60.5; 1.06] showed broader neuronal expression with enrichment in cortical neurons. | Well-supported cortical excitatory population characterized by the combined enrichment of miR-128-3p and miR-132-3p. miR-24-3p provided additional, but less cell-type-specific, cortical support. |
| **Cortico-hippocampal excitatory neurons (n = 5,606)** | Excitatory and hippocampal support, including Slc17a7, Bdnf, Cpne6, Cpne4, Shisa6, Cabp7, Nrn1 and Epha4. | No highly specific miRNA was identified. miR-138-5p [78.3; 1.72] and miR-129-2-3p [49.8; 1.04] supported neuronal identity but were shared across populations. miR-324-5p [17.9; 0.67] showed weaker regional enrichment. | The population was supported primarily by its mRNA profile. The associated miRNAs were broadly neuronal and did not uniquely distinguish the cortico-hippocampal population. |
| **Lateral cortical/piriform neurons (n = 12,312)** | Cortical excitatory support, including Slc17a7, Neurod6, Satb2, Nrn1, Cpne6 and Epha4. Piriform regional identity was supported predominantly by spatial localization. | miR-128-3p [92.0; 1.75], miR-138-5p [93.9; 1.75] and miR-132-3p [73.7; 1.66] showed strong but shared cortical or neuronal enrichment. miR-182 [0.66; 1.25] showed low abundance but reproducible detection in piriform regions across tissue sections. | The population displayed a lateral cortical and piriform spatial distribution, but its miRNA profile was dominated by markers shared with other neuronal populations. Regional annotation therefore relied on combined mRNA and spatial evidence. |
| **GABAergic inhibitory neurons (n = 6,248)** | Strong inhibitory neuronal support, including Gad1, Gad2, Pvalb, Cdh13, Cntnap4, Kcnmb2 and Chrm2. | miR-127-3p [73.6; 2.14] represented the principal associated miRNA but was not exclusive to this population. miR-382-5p [20.7; 1.24] provided additional support and was shared with ventral and subcortical populations. miR-133a-3p [2.2; log2FC not reported] showed low quantitative enrichment but localized to neurons in the thalamic reticular nucleus. | Inhibitory neuronal identity was strongly supported by the mRNA profile. The miRNA signature was supportive but not population-specific, with miR-127-3p showing the strongest association. |
| **Ventral diencephalic neurons (n = 7,903)** | Subcortical and inhibitory neuronal support, including Gad2, Calb2, Cdh13, Cntnap4, Dner, Cacna2d2, Rims3 and Nrp2. | miR-495-3p [50.1; 2.12] and miR-335-5p [53.1; 1.70]. | High-confidence regional neuronal population supported by concordant mRNA expression, spatial localization and enrichment of miR-495-3p and miR-335-5p. |
| **Thalamic-subcortical mixed neurons (n = 9,443)** | Mixed subcortical neuronal support, including Slc17a6, Dner, Cpne6, Cntn6, Cntnap4, Necab2 and Calb2, consistent with a heterogeneous cellular composition. | No population-specific miRNA was identified. miR-129-2-3p [89.1; 1.42] and miR-138-5p [53.9; 1.00] showed strong but broadly neuronal enrichment. | Heterogeneous thalamic and subcortical population characterized by shared neuronal miRNAs rather than a distinct population-specific miRNA signature. |
| **Striatal medium spiny neurons (n = 10,423)** | Strong striatal medium spiny neuron support, including Bcl11b, Ppp1r1b, Meis2, Penk, Pde7b, Wfs1 and Gucy1a1. | miR-139-5p [148.9; 3.27] and miR-153-3p [59.0; 1.58]. | High-confidence striatal population supported by concordant mRNA and miRNA profiles. miR-139-5p showed particularly strong and spatially restricted enrichment. |

| Cell population (n cells) | mRNA support | Primary miRNA marker(s) [score; log2FC] | Integrated conclusion |
|---|---|---|---|
| **Mature oligodendrocytes (n = 12,489)** | Strong oligodendroglial support, including Sox10, Opalin, Gjc3, Cdh20, Gng12 and Tmem163, consistent with mature white-matter oligodendrocytes. | miR-338-3p [151.7; 5.03]. miR-100-5p [98.5; 2.36] and miR-151-5p [77.1; 2.62] showed strong but less specific enrichment. | High-confidence mature oligodendrocyte population. miR-338-3p represented the principal marker, supported by concordant quantitative and spatial evidence. |
| **Choroid plexus/ependymal cells (n = 2,784)** | Strong ventricular and choroid plexus support, including Trp73, Spag16, Slc13a4, Igf2, Cd24a, Paqr5 and Slit2. The combined annotation may encompass two closely related populations. | miR-34b-5p [84.3; 7.95], miR-204-5p [60.4; 5.15], miR-34c-5p [46.4; 7.67] and miR-1298-5p [45.9; 8.02]. | High-confidence ventricular and choroid plexus-associated module defined by a multi-miRNA signature that strongly distinguished this compartment from other populations. |
| **Vascular mural cells (n = 7,681)** | Strong mural and perivascular support, including Carmn, Cspg4, Acta2, Ccn2, Ano1, Dcn and collagen-associated extracellular matrix genes. Endothelial marker expression suggested close spatial association or partial vascular admixture. | miR-143-3p [143.8; 5.34] and miR-145a-5p [69.5; 5.45]. | High-confidence mural-cell population characterized by coordinated enrichment of miR-143-3p and miR-145a-5p, representing one of the clearest miRNA modules in the dataset. |
| **Glial/vascular-associated cells (n = 5,963)** | Mixed molecular support. Gfap, Pdgfra, Sox10, Gjc3 and Cdh20 indicated glial components, whereas Col1a1, Fmod and Aldh1a2 supported stromal or perivascular components. | No population-specific high-confidence miRNA was identified. | Heterogeneous glial and perivascular compartment rather than a discrete cell type. The mRNA and spatial profiles supported a mixed identity without a specific defining miRNA signature. |

**Supplementary Table 6 | Integrated interpretation of miRNA-defined cell populations and associated miRNA signatures**. The table summarizes the 12 annotated cell populations, including cell number, supporting mRNA markers, primary enriched miRNAs with their corresponding scores and log2 fold changes, and an integrated molecular and spatial interpretation. High-confidence populations are marked in the table and were defined by concordant mRNA identity, miRNA enrichment and anatomical localization. Populations lacking a specific miRNA signature were supported primarily by shared neuronal or glial miRNA profiles together with mRNA expression and spatial distribution.

#### Supplementary Table 7. Single-cell assessment of miRNA markers

Supplementary Table 7 provides an integrated anatomical and single-cell assessment of the miRNA markers evaluated in this study. For each miRNA, the table reports the strength of the evidence, its principal regional and cellular association, the corresponding Wilcoxon score and fold-change value, and an integrated interpretation combining spatial localization with cell-population enrichment. The analysis identifies high-confidence markers for endothelial cells, vascular mural cells, mature oligodendrocytes, striatal medium spiny neurons, ventral diencephalic neurons, and choroid plexus/ependymal populations. Other miRNAs showed broader neuronal, glial, vascular, or subcortical distributions and were therefore classified as shared or supporting markers rather than population-specific markers. miRNAs lacking sufficient quantitative or spatial support were classified as incomplete, weak, or unsupported associations.

| miRNA | Assessment | Regional and cellular association | Wilcoxon statistics | Integrated anatomical and single-cell analysis |
|---|---|---|---|---|
| **miR-24-3p** | Shared marker | Broad neuronal expression with cortical enrichment; associated with cortico-hippocampal excitatory neurons. | S=60.51; FC=1.06 | Single-cell enrichment was consistent with the anatomical distribution. Shared across excitatory populations; not specific. |
| **miR-132-3p** | Good cell-type marker | Cortical enrichment; associated with cortical excitatory neurons. | S=86.12; FC=1.74 | Anatomical localization and single-cell enrichment were concordant. Strong cortical association, but not exclusive. |
| **miR-127-3p** | Shared marker | Broad neuronal expression with ventral enrichment; associated with inhibitory and subcortical neurons. | S=73.60; FC=2.14 | Single-cell enrichment was consistent with the anatomical distribution. Supports inhibitory identity, but is shared subcortically. |
| **let-7e-5p** | Shared marker | Broad neuronal expression; associated with inhibitory and subcortical neurons. | S=62.49; FC=1.12 | Single-cell enrichment was consistent with the anatomical distribution. Shared inhibitory-subcortical signal. |
| **miR-107-3p** | Shared marker | Broad neuronal expression, including hippocampal and olfactory territories; associated with inhibitory neurons. | S=49.42; FC=1.23 | Anatomical localization and single-cell enrichment were concordant. Broad inhibitory/olfactory association. |
| **miR-128-3p** | Good cell-type marker | Cortical enrichment; associated with cortical and cortico-hippocampal excitatory neurons. | S=123.15; FC=2.04 | Anatomical localization and single-cell enrichment were concordant. Strong cortical marker, although not fully exclusive. |
| **miR-133a-3p** | Insufficient evidence | Inhibitory-neuron and internal-capsule enrichment; associated with inhibitory neurons. | S=2.25; FC=1.68 | The anatomical distribution is compatible with the proposed association, but single-cell Wilcoxon support is weak. |
| **miR-301a-3p** | Insufficient evidence | Broad neuronal expression; associated with mixed neuronal and glial populations. | S=ND; FC=ND | The anatomical signal alone is insufficient for robust assignment in the absence of differential-expression statistics. |
| **miR-370-3p** | Insufficient evidence | Broad neuronal expression; associated with neuronal populations. | S=ND; FC=ND | The anatomical distribution appears broadly neuronal and poorly specific; single-cell differential-expression support is unavailable. |
| **miR-379-5p** | Insufficient evidence | Broad neuronal expression; associated with neuronal populations. | S=ND; FC=ND | The combined anatomical and single-cell pattern is consistent with a shared neuronal signal rather than a population-specific marker. |
| **miR-485-5p** | Insufficient evidence | Broad neuronal expression; associated with subcortical and inhibitory neurons. | S=ND; FC=ND | The anatomical distribution is compatible with a subcortical or inhibitory association, but single-cell Wilcoxon evidence is unavailable. |
| **miR-143-3p** | Good cell-type marker | Vascular mural-cell enrichment; associated with vascular mural cells. | S=143.79; FC=5.34 | Anatomical localization and single-cell enrichment were concordant. Strong mural marker; not endothelial-specific. |
| **miR-145a-5p** | Good cell-type marker | Vascular mural-cell enrichment; associated with vascular mural cells. | S=69.52; FC=5.45 | Anatomical localization and single-cell enrichment were concordant. Highly consistent mural-cell marker. |
| **miR-195a-5p** | Insufficient evidence | Cerebellar enrichment; associated with vascular mural cells in the single-cell analysis. | S=ND; FC=ND | The anatomical distribution is compatible with a secondary mural association, but single-cell quantitative evidence is incomplete. |

| miRNA | Assessment | Regional and cellular association | Wilcoxon statistics | Integrated anatomical and single-cell analysis |
|---|---|---|---|---|
| **miR-152-3p** | Insufficient evidence | Meningeal-surface enrichment; associated with mural and endothelial compartments. | S=ND; FC=ND | The anatomical distribution is compatible with vascular enrichment, but single-cell evidence is incomplete. |
| **miR-126a-3p** | Good cell-type marker | Endothelial and vascular enrichment; associated with endothelial cells. | S=151.44; FC=3.87 | Strongest endothelial-associated signal; minor shared vascular expression remains. |
| **miR-150-5p** | Good cell-type marker | Endothelial and vascular enrichment; associated with endothelial cells. | S=117.23; FC=3.83 | Single-cell enrichment was concordant with the anatomical distribution. Strong endothelial marker supported by Wilcoxon. |
| **miR-338-3p** | Good cell-type marker | White-matter and mature-oligodendrocyte enrichment; associated with mature oligodendrocytes. | S=151.65; FC=5.03 | Anatomical localization and single-cell enrichment were concordant. Most robust mature-oligodendrocyte marker. |
| **miR-151-5p** | Good supporting marker | Broad neuronal and cerebrovascular expression; associated with mature oligodendrocytes and fiber tracts. | S=77.06; FC=2.62 | Anatomical localization and single-cell enrichment were concordant. Strong oligodendrocyte/tract support marker. |
| **miR-21a-5p** | Shared marker | Broad expression with white-matter and fiber-tract enrichment; associated with oligodendrocytes. | S=77.66; FC=2.73 | Supports oligodendroglial/tract identity, but is not specific. |
| **miR-100-5p** | Good supporting marker | White-matter and mature-oligodendrocyte enrichment; associated with mature oligodendrocytes. | S=98.55; FC=2.36 | Good oligodendrocyte-associated marker, with some expression elsewhere. |
| **miR-139-5p** | Good regional marker | Striatal and medium-spiny-neuron enrichment; associated with striatal medium spiny neurons. | S=148.89; FC=3.27 | Anatomical localization and single-cell enrichment were concordant. One of the strongest and most regionally restricted striatal markers. |
| **miR-138-5p** | Shared marker | Broad neuronal expression with hippocampal and striatal enrichment; associated with striatal and cortical neuronal populations. | S=121.79; FC=2.05 | Strong neuronal miRNA with striatal bias, but not population-specific. |
| **miR-153-3p** | Good cell-type marker | Striatal and medium-spiny-neuron enrichment; associated with striatal medium spiny neurons. | S=58.99; FC=1.58 | Supported striatal-associated marker, although not completely exclusive. |
| **miR-218-5p** | Supporting regional marker | Broad neuronal expression with hippocampal and striatal enrichment; associated with cortical and hippocampal neuronal populations. | S=18.05; FC=1.20 | Anatomical localization supports a cortical–hippocampal association, but single-cell Wilcoxon support is modest. |
| **miR-344-3p** | Incomplete evidence | Broad neuronal expression with striatal enrichment; associated with subcortical neuronal populations. | S=ND; FC=ND | The anatomical distribution suggests regional structure, but differential-expression statistics are unavailable. |
| **miR-409-3p** | Weak evidence | Broad neuronal expression; associated with cortico-hippocampal excitatory neurons. | S=3.24; FC=0.76 | The anatomical direction is plausible, but the single-cell Wilcoxon score is too low to support marker status. |

| miRNA | Assessment | Regional and cellular association | Wilcoxon statistics | Integrated anatomical and single-cell analysis |
|---|---|---|---|---|
| **miR-129-2-3p** | Shared regional marker | Broad neuronal expression with reduced signal in fiber tracts; associated with diencephalic and subcortical neurons. | S=99.09; FC=1.72 | Defines a broad diencephalic-subcortical neuronal program; not cell-population-specific. |
| **miR-495-3p** | Good cell-type marker | Ventral diencephalic enrichment; associated with ventral diencephalic neurons. | S=50.07; FC=2.12 | Anatomical localization and single-cell enrichment were concordant. Strong ventral-diencephalic marker. |
| **miR-335-5p** | Good cell-type marker | Broad neuronal expression with ventral enrichment; associated with ventral diencephalic neurons. | S=53.06; FC=1.70 | Anatomical localization and single-cell enrichment were concordant. Good regional marker, although not fully specific. |
| **miR-382-5p** | Shared regional marker | Broad expression with ventral enrichment; associated with ventral diencephalic and inhibitory neurons. | S=35.32; FC=1.82 | Supports a shared ventral-subcortical/inhibitory module rather than one population. |
| **miR-381-3p** | Shared regional marker | Broad expression with ventral enrichment; associated with ventral diencephalic and striatal neurons. | S=35.89; FC=1.35 | Shared subcortical marker with limited cell-population specificity. |
| **miR-487b-3p** | Supporting regional marker | Broad expression with ventral enrichment; associated with ventral diencephalic neurons. | S=23.57; FC=1.29 | Secondary ventral-diencephalic association supported by the integrated analysis. |
| **miR-7a-5p** | Incomplete evidence | Hypothalamic enrichment; associated with hypothalamic and other neuronal populations. | S=ND; FC=ND | The anatomical distribution suggests septal enrichment, but statistical support is unavailable. |
| **miR-324-5p** | Supporting marker | Broad neuronal expression with striatal enrichment; associated with cortico-hippocampal excitatory neurons. | S=17.95; FC=0.67 | Moderate single-cell Wilcoxon support and compatible anatomical localization indicate a secondary rather than defining marker. |
| **miR-467a** | Incomplete evidence | Broad neuronal expression; associated with diencephalic and subcortical neurons. | S=ND; FC=ND | The anatomical distribution is compatible with a regional association, but single-cell Wilcoxon statistics are unavailable. |
| **miR-34b-5p** | Good cell-type marker | Choroid-plexus and ventricular enrichment; associated with choroid plexus and ependymal cells. | S=84.29; FC=7.95 | Anatomical localization and single-cell enrichment were concordant. Strong ventricular/choroid-plexus marker. |
| **miR-1298-5p** | Good cell-type marker | Choroid-plexus and ventricular enrichment; associated with choroid plexus and ependymal cells. | S=45.87; FC=8.02 | Anatomical localization and single-cell enrichment were concordant. Highly specific ventricular/choroid-plexus marker. |
| **miR-34c-5p** | Good cell-type marker | Ventricular-system enrichment; associated with choroid plexus and ependymal cells. | S=46.43; FC=7.67 | Anatomical localization and single-cell enrichment were concordant. Highly consistent ventricular marker. |
| **miR-204-5p** | Good cell-type marker | Choroid-plexus and ventricular enrichment; associated with choroid plexus and ependymal cells. | S=60.35; FC=5.15 | Strong ventricular marker; should not be assigned primarily to mixed glial/vascular cells. |

| miRNA | Assessment | Regional and cellular association | Wilcoxon statistics | Integrated anatomical and single-cell analysis |
|---|---|---|---|---|
| **miR-130a-3p** | Shared marker | Broad neuronal expression with ventral enrichment; associated with glial, vascular and mural compartments. | S=50.10; FC=1.88 | Shared vascular/perivascular marker; not population-specific. |
| **miR-34b-3p** | Incomplete evidence | Choroid-plexus and ventricular enrichment; associated with choroid plexus and ependymal cells. | S=ND; FC=ND | The anatomical distribution is compatible with the ventricular miR-34 module, but single-cell quantitative evidence is incomplete. |
| **miR-140-3p** | Insufficient evidence | Broad neuronal expression; associated with glial, vascular and mural compartments. | S=ND; FC=ND | The anatomical distribution is compatible with vascular enrichment, but single-cell quantitative evidence is unavailable. |
| **miR-182-5p** | Not supported | Piriform-neuron enrichment; associated with lateral cortical and piriform neurons. | S=0.66; FC=1.25 | Anatomical localization alone does not support marker assignment because the single-cell Wilcoxon result is not significant. |
| **miR-19b-3p** | Shared marker | Broad neuronal expression; associated with glial and vascular-associated cells. | S=22.32; FC=0.90 | Supports a mixed glial/vascular signal, but lacks strong specificity. |
| **miR-99a-5p** | Shared marker | Broad neuronal expression; associated with glial, vascular and hippocampal compartments. | S=43.81; FC=1.89 | Useful regional association, but shared across compartments. |
| **miR-30c-5p** | Shared marker | Broad neuronal expression, including the SVZ, olfactory bulb and hippocampus; associated with inhibitory and glial/vascular populations. | S=42.27; FC=0.77 | Broad marker with limited specificity. |
| **miR-141-3p** | Insufficient evidence | Olfactory-bulb enrichment; associated with glial and vascular-associated cells. | S=ND; FC=ND | The combined anatomical and single-cell evidence is insufficient for marker assignment. |
| **miR-26b-5p** | Weak evidence | Broad neuronal expression; associated with mature oligodendrocytes and other glial cells. | S=2.09; FC=0.48 | Glial direction is plausible, but score and effect size are too low. |

**Supplementary Table 7 | Single-cell assessment of miRNA markers**. For each miRNA, the table summarizes the level of evidence, regional and cellular association, Wilcoxon score and fold-change value, and an integrated interpretation combining anatomical localization with single-cell enrichment. Markers with concordant spatial, quantitative and cell-population evidence were classified as high-confidence or supporting markers, whereas broadly distributed miRNAs were designated as shared markers. Associations lacking sufficient statistical or spatial support were classified as weak, incomplete or unsupported.

#### Supplementary Table 8. Best-supported miRNA markers of high-confidence cell populations

Supplementary Table 8 summarizes the best-supported miRNA markers associated with the high-confidence cell populations identified in the mouse brain. For each population, the table reports the selected miRNA markers, their Wilcoxon scores, and an integrated interpretation based on differential enrichment, anatomical localization, and concordance with single-cell identities. The strongest and most specific associations were observed for endothelial cells, vascular mural cells, mature oligodendrocytes, striatal medium spiny neurons, and choroid plexus/ependymal cells, whereas cortical excitatory and ventral diencephalic populations were defined by regionally informative combinations of miRNA markers. Supporting miRNAs with broader spatial distributions are indicated where they reinforce, but do not uniquely define, the corresponding cell population.

| Cell population | Best-supported miRNA marker(s) | Wilcoxon score | Integrated anatomical and single-cell interpretation |
|---|---|---|---|
| **Endothelial cells** | miR-150-5p; miR-126a-3p | 117.2; 151.4 | miR-150-5p showed the more endothelial-restricted association, whereas miR-126a-3p displayed strong but broader vascular enrichment and was also detected in mural cells. |
| **Cortical excitatory neurons** | miR-128-3p; miR-132-3p | 123.2; 86.1 | These miRNAs represented the best-supported cortical excitatory markers. Both showed concordant cortical localization, although expression was shared with related cortical neuronal populations. |
| **Ventral diencephalic neurons** | miR-495-3p; miR-335-5p | 50.1; 53.1 | Both miRNAs provided regionally informative signatures, with stronger enrichment in ventral diencephalic neurons than in the other neuronal populations analysed. |
| **Striatal medium spiny neurons** | miR-139-5p; miR-153-3p | 148.9; 59.0 | Both miRNAs showed strong striatal enrichment. miR-139-5p displayed particularly high specificity and clear separation from other populations. |
| **Mature oligodendrocytes** | miR-338-3p; miR-151-5p | 151.7; 77.1 | Both miRNAs supported an oligodendroglial identity. miR-338-3p was among the most specific markers detected in the dataset. |
| **Choroid plexus / ependymal cells** | miR-34b-5p; miR-204-5p; miR-34c-5p; miR-1298-5p | 84.3; 60.4; 46.4; 45.9 | The combined signature defined a distinct ventricular and choroid plexus-associated module with strong spatial and differential-expression concordance. |
| **Vascular mural cells** | miR-143-3p; miR-145a-5p | 143.8; 69.5 | Both miRNAs showed strong mural-cell enrichment. miR-143-3p was therefore interpreted as a mural-cell marker rather than an endothelial-specific marker. |

**Supplementary Table 8 | Best-supported miRNA markers of high-confidence cell populations**. The table lists the miRNA markers associated with the high-confidence cell populations identified in the mouse brain, together with their Wilcoxon scores and an integrated interpretation based on differential enrichment, anatomical localization and concordance with single-cell identities. Primary markers are distinguished from supporting miRNAs with broader regional or cellular distributions.

**Supplementary Table 6. Tissue sections used in the development of miR-Space**

Supplementary Table 6 summarizes the tissue sections used during the development, optimization and validation of miR-Space. For each section, the table provides the experimental identifier, a concise description of the experimental approach and its purpose, the assay plex and the platform used. Experimental identifiers follow the format S-M/H-B-V-C, where S denotes the section number, M the mouse used, H the human sample used, B the experimental batch, V the barcode version and C the codebook used for decoding. The dataset includes workflow comparisons, probe and linker controls, ligation and barcode evaluations, reproducibility experiments, human-tissue applications, high-plex miRNA-mRNA co-detection assays and spatial single-cell analyses performed on the Xenium platform. Together, these experiments document the technical progression of miR-Space from initial assay optimization to high-plex and spatially resolved validation.

| Code / Identifier | Experimental approach | Assay plex | Platform |
|---|---|---|---|
| **S1** M1_S1_B1_V1_C1_Standard | Initial tissue section used to evaluate barcode version 1 of miR-Space. | 50-plex | In-house |
| **S2** M1_S2_B1_V1_C1_Duplex | Pre-hybridization of Link A and Link B before tissue application to test whether probe pairing improved miRNA detection. | 50-plex | In-house |
| **S3** M1_S3_B1_V1_C1_USER_Standard | USER-mediated removal of Link A to evaluate enzymatic elimination. | 50-plex | In-house |
| **S4** M1_S4_B1_V1_C1_USER_Duplex | Combined Link pre-hybridization and USER-mediated probe removal to assess whether both workflow modifications improved detection. | 50-plex | In-house |
| **S5** M2_S5_B2_V1_C1_Second_Positive | Technical replicate of the initial experiment using barcode version 1. | 50-plex | In-house |
| **S6** M2_S6_B2_V2_C1_Standard | First evaluation of barcode version 2 to assess a new barcode design. | 50-plex | In-house |
| **S7** M2_S7_B2_V1_C1_Direct_PLPs | Direct application of PLPs without miRNA extension to test the feasibility of direct detection. | 50-plex | In-house |
| **S8** M2_S8_B2_V1_C1_Wrong_Match | Incorrect pairing of Link A and Link B to assess assay specificity and nonspecific background. | No-Extension control | In-house |
| **S9** M2_S9_B2_V1_C1_OnlyPLPs | Application of PLPs targeting extended miRNAs in the absence of the linkers required for the extension reaction. | No-Extension control | In-house |
| **S10** M2_S10_B2_V1_C1_Short_Linker | Application of PLPs targeting extended miRNAs in the absence of the Linker A required for the extension reaction. | No-Extension control | In-house |
| **S11** M3_S11_B3_V2_C2_C3_50miRNA_50mRNA | Joint spatial detection of 50 miRNAs and 50 mRNAs in the same tissue section to demonstrate simultaneous analysis of both RNA classes. | 100-plex | In-house |
| **S12** M3_S12_B4_V2_C4_200miRNA_50mRNA | High-plex co-detection of 200 miRNAs and 50 mRNAs to evaluate scaling of simultaneous miRNA and mRNA detection. | 250-plex | In-house |
| **S13** H1_S13_B5_V2_C5_Human_Positive_1 | Spatial miRNA detection in a first human tissue section to extend the method from mouse to human samples. | 50-plex | In-house |
| **S14** H1_S14_B5_V2_C5_Human_Positive_2 | Technical replicate in a consecutive human sample. | 50-plex | In-house |
| **S15** H1_S15_B5_V2_C5_Human_Negative | Matched human no-extension control in the absence of the Linkers required for the extension reaction. | No-Extension control | In-house |
| **S16** M3_S16_B6_V2_C1_SplintR | SplintR-mediated ligation of PLPs to compare with KOD RNA Ligase. | 50-plex | In-house |
| **S17** M_S17_B6_V2_C1_KOD | Evaluation of KOD RNA Ligase as an alternative ligation enzyme to determine whether it produced similar spatial patterns. | 50-plex | In-house |
| **S18** M3_S18_B7_V2_C1_Second_Positive | Technical replicate of the experiment using barcode version 2. | 50-plex | In-house |

| Code / Identifier | Experimental approach | Assay plex | Platform |
|---|---|---|---|
| **S19** M4_S19_B8_V3_C6_Sagittal | Barcode V3 tested in a sagittal mouse section to support transition to the next barcode generation. | 50-plex | In-house |
| **S20** M5_S20_B8_V3_C6_Positive | Barcode version 3 was tested in a coronal mouse brain section using the same panel as the Xenium experiments to assess cross-platform reproducibility. | 50-plex | In-house |
| **S21** M5_S21_B9_V3_C6_Negative | Matched V3 no-extension control in the absence of the Linkers required for the extension reaction. | No-Extension control | In-house |
| **S22** M5_S22_B9_V3_C8_200miRNA_50mRNA | Co-detection of 200 miRNAs and 50 mRNAs to explore novel miRNAs in the mouse brain. | 250-plex | In-house |
| **S23** M5_S23_B9_V3_C8_Negative | Matched negative control for the exploratory 250-plex assay in the absence of the Linkers required for the extension reaction. | No-Extension control | In-house |
| **S24** M5_S24_B10__250plex_1 | Technical replicate of the 250-plex tissue experiment using barcode version 2. | 250-plex | In-house |
| **S25** M6_S25_B10__250plex_2 | Technical replicate of the 250-plex tissue experiment using barcode version 2. | 250-plex | In-house |
| **S26** M7_S26_B11_V3_C9_BL6_adult1 | Coronal adult mouse section analysed by Xenium for spatial single-cell integration of miRNA detection and transcriptomics. | 297-plex | Xenium |
| **S27** M7_S27_B11_V3_C9_BL6_adult2 | Consecutive coronal adult mouse section analysed by Xenium to provide an additional biological sample for comparison. | 297-plex | Xenium |
| **S28** M8_S28_B11_V3_C9_P28_coronal | Coronal P28 mouse section analysed by Xenium to extend spatial single-cell analysis to a younger developmental stage. | 297-plex | Xenium |
| **S29** M9_S29_B11_V3_C9_P16_coronal | Coronal P16 mouse section analysed by Xenium to add an earlier developmental stage for spatial comparison. | 297-plex | Xenium |
| **S30** M10_S30_B11_V3_C9_sagittal_adult | Sagittal P16 mouse section analysed by Xenium to compare tissue orientations in spatial single-cell analysis. | 297-plex | Xenium |

**Supplementary Table 6 | Tissue sections used in the development of miR-Space.** The table lists the experimental identifier, summarized experimental approach, assay plex and platform for each tissue section used during miR-Space development, optimization and validation. S denotes the section number, M the mouse used, H the human sample used, B the experimental batch, V the barcode version and C the codebook used for decoding. The experiments comprise workflow optimization, technical controls, reproducibility analyses, applications to human tissue, high-plex miRNA–mRNA co-detection and Xenium-based spatial single-cell profiling.
